# *In silico* subtractive genomics and reverse vaccinology approach reveal outer membrane proteins OMP47 and LptD as dual-purpose drug and vaccine targets in *Pasteurella multocida* Pm70

**DOI:** 10.64898/2026.10.07.757254

**Authors:** Ashiqur Rahman Khan Chowdhury, Sharita Islam, Asma Farid, Farzia Lamia, Tasmim Fahima Ahmad, Neyma Jahan Hossain, Aurpita Rani Paul, Tabassum Afrin, Sabrina Moriom Elias

## Abstract

Drug targets and vaccines for bacterial diseases are challenging to develop. Computational subtractive genomics and reverse vaccinology offer a viable substitute for conventional drug and vaccine development approaches. In recent years, *Pasteurella multocida,* a Gram-negative pathogen responsible for severe infections in humans along with major diseases in livestock and poultry, has developed resistance to multiple drugs. As a result, there is an urgent demand for research into reliable, efficient, highly potent, and cost-effective strategies to combat the rise of *P. multocida* infections. To address this, the present study integrated an *in silico* subtractive genomics and reverse vaccinology workflow to pinpoint proteins with potential value as both drug and vaccine targets. From the complete *P. multocida* Pm70 proteome, we identified 446 proteins that had no similarities with human proteins, out of which 129 were found to be essential for the pathogen’s survival. These 129 proteins are therefore potential candidates for drug development. Further metabolic pathway analysis, virulence potential, subcellular localization, and structural analyses revealed two outer membrane proteins, namely, 47 kDa outer membrane protein and lipopolysaccharide-assembly protein LptD, as promising candidates for therapeutic development. Subsequent antigenicity analysis revealed both outer membrane proteins as potential vaccine targets, exhibiting highly favorable antigenicity scores. Candidate B– and T-cell epitopes were systematically mapped from these two targets based on antigenicity, immunogenicity, toxicity, and allergenicity potential and only high-confidence epitopes were assembled using adjuvants and class-specific linkers to construct the final multi-epitope vaccine (MEV). The designed MEV construct was demonstrated to be highly antigenic, non-toxic and non-allergenic to humans, while possessing favorable structural and physicochemical properties, and exhibited strong binding interactions with human toll-like receptors with minimal molecular deformability. Although further experimental validation is recommended to fully assess their efficacy, the present study establishes a solid foundation for the development of novel therapeutics and vaccine candidates against *P. multocida*, ultimately contributing to the reduction of its rising infections.

## Introduction

*Pasteurella multocida* is a zoonotic Gram-negative pathogen that infects a wide range of hosts, including cattle, buffalo, sheep, goats, pigs, rabbits, poultry, as well as humans [1–7]. It is responsible for severe diseases such as haemorrhagic septicaemia and bovine respiratory disease in ruminants, pneumonic pasteurellosis in swine, pasteurellosis in rabbits, and fowl cholera in birds [1–5]. It also causes severe infections in immunocompromised or otherwise high-risk human populations [3,6,7]. Because of its broad host range, and its ability to establish both acute and persistent infections, *P. multocida* remains a substantial worldwide burden to human and animal health.

*P. multocida* infection is frequently linked to exposure to colonized companion animals, particularly cats and dogs, which carry the bacterium in their oral flora at high rates, approximately 70 to 90% in cats and 20 to 50% in dogs [3,8]. Transmission usually follows bites, scratches, or saliva contamination of broken skin [3,8]. *Pasteurella* species, predominantly *P. multocida*, are recovered from about 75% of infected cat-bite wounds and about 50% of infected dog-bite wounds [3,8–10]. In the United States alone, animal bites account for roughly 4-5 million injuries and about 300,000 emergency-department visits annually [3,7]. Infections such as skin and soft-tissue infection can develop rapidly, and respiratory and invasive disease, such as tracheobronchitis, pneumonia, bacteremia, urinary tract infection, empyema, lung abscesses, meningitis, septic arthritis, and peritonitis can also occur, particularly in older adults and in patients with chronic liver disease, diabetes, chronic lung disease, malignancy, or immunosuppression [3,8]. Respiratory infection typically exhibits symptoms such as fever, cough, chest pain, and dyspnea [3,9,11–13]. Severe invasive disease has mortality rates of approximately 25 to 30% [3,9,11–13].

The management of *P. multocida* infections is increasingly challenged by antimicrobial resistance [14,15]. Clinical and veterinary isolates have been reported with increased resistance to several antibiotics including β-lactams, tetracyclines, macrolides, and fluoroquinolones [14,15]. Moreover, in human cases, resistance to erythromycin, fluoroquinolones, amoxicillin-clavulanate, penicillin, and other β-lactams have been observed, despite penicillin being the recommended first-line of treatment [7,16,17]. These challenges are often linked to extensive antimicrobial use in animal husbandry as well as in human clinical practice [18,19]. It is important to note that existing vaccines remain largely confined to veterinary use and no approved human vaccine is available [20–23]. Hence the emergence of multidrug-resistant strains highlights the need for alternative treatment options [7,14,15].

The significant advancements in computational biology and bioinformatics have revolutionized the field of drug design, leading to lower costs and faster timelines compared to traditional laboratory trials [24]. One of the main strategies employed is the subtractive genomics approach which enables the prioritization of pathogen-specific proteins that are essential for the survival of the pathogen by excluding those having similarities with host proteins thereby reducing the risk of cross-reactivity with the host [25–29]. In parallel, reverse vaccinology extends this framework by identifying antigenic proteins, recognizing immunogenic high confidence epitopes, and assembling them into a multi-epitope vaccine construct [30–33]. These techniques have been proven to be valuable in uncovering potential treatment targets in a wide variety of pathogenic strains [26–47]. Previous studies have investigated drug targets through subtractive genomics and vaccine candidates through outer-membrane protein characterization and epitope prediction in *P. multocida* [48–51]. Building on these approaches, this study evaluates OMP47 and LptD jointly as candidate drug targets and antigen sources for a human-oriented vaccine construct.

In the present study, the integrative subtractive genomics and reverse vaccinology pipeline was applied to the complete *P. multocida* Pm70 proteome to recognize proteins with potential value as both drug and vaccine targets. Sequential filtering based on host similarity, essentiality in pathogen survival, involvement in important metabolic pathways, druggability investigations, virulence association, subcellular localization, protein network analysis, and structural investigations prioritized two promising target proteins, namely 47 kDa outer membrane protein and LPS-assembly protein LptD. The 47 kDa outer membrane protein, or OMP47, functions in adhesion and transport in the outer membrane region of the pathogen, whereas, LptD is involved in inserting lipopolysaccharide (LPS) molecules into the outer membrane’s outer leaflet contributing to the pathogen’s outer membrane assembly, permeability barrier function, and structural stability [52–64]. Furthermore, the antigenic potential of these two proteins was evaluated and high-confidence epitopes with immunogenic potential were selected. These epitopes were then concatenated into a single MEV construct using adjuvants and linkers, and the resulting MEV was assessed for physiochemical and structural properties, its engagement with TLR4, its capability for immune response, and its suitability for recombinant expression in bacterial systems. Although further experimental validation is imperative, we anticipate that the prioritized outer membrane proteins and the proposed vaccine construct will provide a solid framework for future drug and vaccine development against infections caused by *P. multocida*.

## Materials and Methods

To identify proteins as potential drug targets and to design a MEV candidate targeting *P. multocida*, the subtractive genomics workflow followed by the reverse vaccinology pipeline was applied, as illustrated in **Fig 1**.

**Fig 1.**
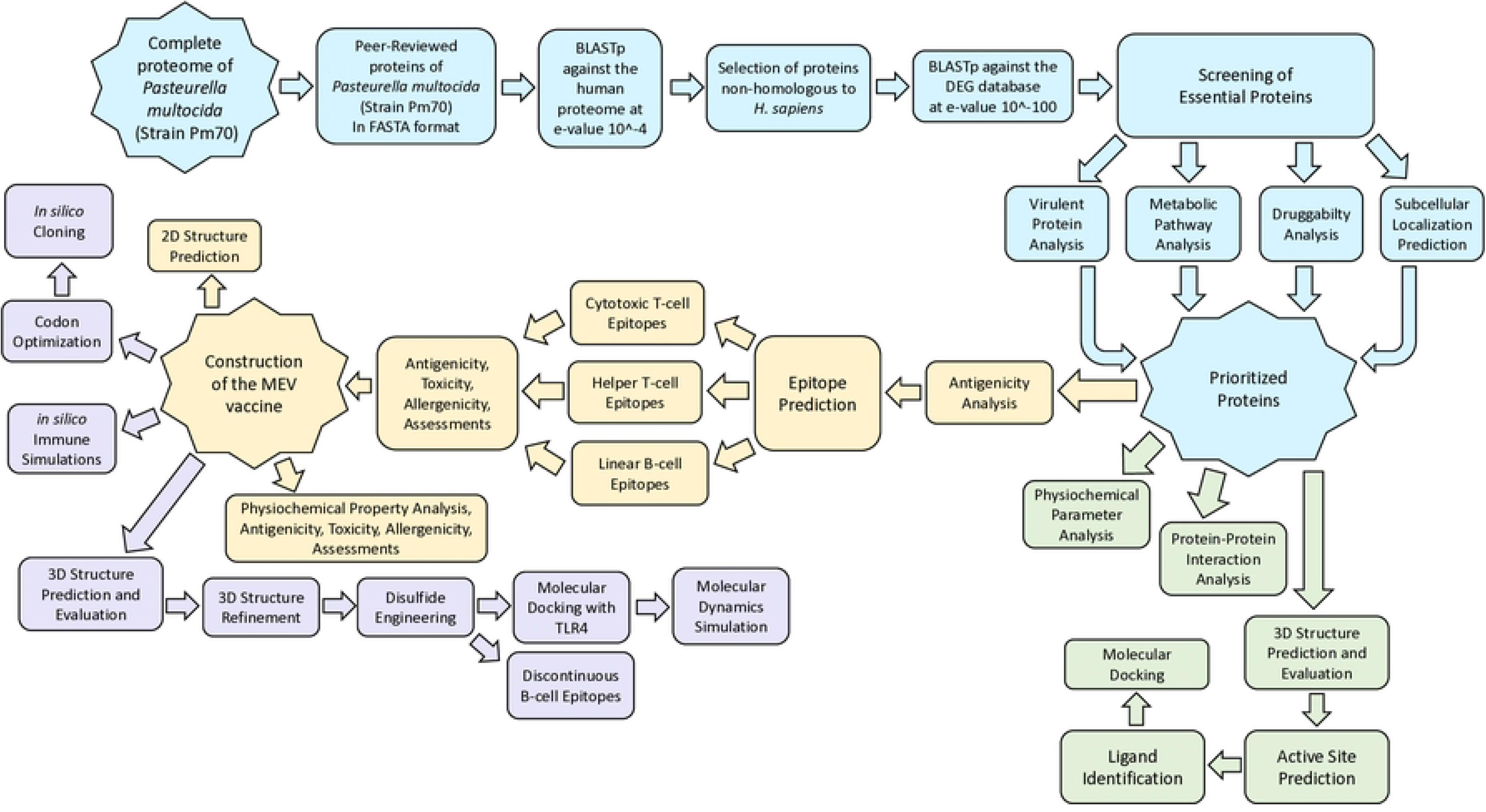
Schematic flowchart illustrating the workflow for drug target identification and multi-epitope vaccine design in *P. multocida* Pm70. Light-blue boxes indicate the subtractive genomics approach including the screening of non-homologous and essential proteins. Light-green boxes represent the various structural analyses used in this study. Yellow boxes indicate antigenicity and epitope-based MEV design. Lavender boxes represent the optimization and validation of the constructed MEV. Arrows indicate the direction of the workflow.

### Retrieval of protein sequences and screening of non-homologous and essential proteins

The entire *P. multocida* Pm70 proteome along with its peer-reviewed sequences were retrieved from the UniProt database in FASTA format [65–67]. To identify proteins that have little to no similarities with host sequences, the selected peer-reviewed proteins were subjected to a BLASTp search against the *Homo sapiens* proteome in the NCBI database, keeping the E-value threshold at 10^-4^ [68].

The resulting non-host-homologous proteins were subjected to BLASTp analysis with an E-value cutoff of 10^-100^ against the DEG version 15.2 (Database of Essential Genes) database to recognize proteins that are indispensable for bacterial survival [28,44,69–73]. Proteins demonstrating high sequence similarities with the DEG database entries were considered essential for bacterial viability and were chosen for further analysis [26–28,40,42,44,69,71,73,74].

### Exploration of metabolic pathways, known drug target investigation, virulence association, and subcellular localization

To effectively analyze the involvement of essential proteins in the metabolic pathways of both the host and the pathogen, metabolic pathway analysis was carried out using the KEGG KAAS Server (KEGG Automatic Annotation Server) [75,76]. The organism codes “hsa” for *Homo sapiens* and “pmu” for *P. multocida* were used to retrieve the metabolic pathways of the host and pathogen, respectively. These pathways were then compared against the shortlisted set of essential proteins in order to identify proteins associated with metabolic pathways unique to the pathogen.

The shortlisted proteins were further screened against DrugBank database (version 6.0) using BLASTp with an expectation value threshold of 10^-5^ [77]. Proteins demonstrating significant similarity to database entries were characterized as potentially druggable targets, whereas the ones showing no sequence similarity were classified as novel drug targets.

Virulence potential was then assessed using the VirulentPred 2.0 server, which predicts virulent proteins using machine-learning approaches based on PSSM-derived features [78]. Subsequently, PSORTb v3.0.3 and CELLO v2.5 tools were employed to predict subcellular localization [79–82]. These tools have been proven to be highly effective in accurately anticipating the locations of proteins and classifying them into subcellular compartments, including the outer-membrane, cytoplasm, extracellular space, cytoplasmic membrane, periplasm, and others with unknown origins [79,81,82].

### Physicochemical characterization, protein interaction analysis, and structural modelling

The ExPASy ProtParam tool determined the physicochemical features of the target proteins from their respective amino acid sequences [83]. The protein-protein interaction networks of the target proteins were explored using the STRING database [84]. Only interactions involving nodes connected to ≥3 other proteins and having confidence scores of ≥0.400 were considered, in order to reduce the likelihood of false-positive associations [85].

The SWISS-MODEL server, which applies homology-based modelling to predict tertiary structures from amino-acid sequences, was used to generate the three-dimensional structures of the selected proteins [86]. For each protein, a suitable template was chosen based on the Global Quality Estimate scores (GMQE), sequence identity, and template coverage percentage [87,88]. The obtained models were cross-checked against the AlphaFold Protein Structure Database, an advanced AI system developed by Google DeepMind, which offers an extensive repository with over 214 million entries, effectively providing a comprehensive coverage of the UniProt database [89–91]. Additional structural validation was carried out for the SWISS-MODEL generated structures using ProSA-web and the SAVES v6.1 server, including ERRAT and PROCHECK analyses [92–97].

### Binding-site prediction, ligand identification, and molecular docking

Potential binding sites in the modeled protein structures were identified using the DoGSiteScorer tool of the ProteinPlus server [98,99]. This tool predicts binding-site pockets by analyzing the geometry and physicochemical features of the protein surface [98,99]. With the information on experimentally confirmed ligands being unavailable, to predict likely ligands and evaluate their interactions with the selected targets, we employed the ProBiS server, which utilizes a theoretical approach, employing molecular simulation to predict the most favorable protein ligand [100]. The predicted binding pockets and ligand interactions were visualized using PyMOL v3.1.

Molecular docking between the selected proteins and their predicted ligands was then carried out using PyRx v0.8. The highest-ranked docked complexes were selected for further interpretation, and post-docking interaction analysis and visualization were performed using BIOVIA Discovery Studio v4.0 [101].

### Antigenicity prediction and epitope mapping

To evaluate the antigenicity of the shortlisted outer membrane proteins, the VaxiJen v2.0 server was employed with a ≥0.4 cutoff score [102]. Antigenic outer membrane proteins were then selected for epitope prediction. Utilizing the NetMHCpan EL 4.1 algorithm from the IEDB (Immune Epitope Database) server, the cytotoxic T-cell (CTL) epitopes were identified. Keeping the percentile rank ≤0.4%, prediction scores ≥0.8, commonly occurring HLA-class-I alleles as the reference panel, and peptide length at 9 amino acids as the selection criteria [103]. IEBD MHC I Immunogenicity tool then assessed the CTL candidate epitopes, and the peptides having positive immunogenicity scores were retained [104].

Following that, utilizing the NetMHCIIpan EL 4.1 algorithm from the IEDB server, the helper T-cell (HTL) epitopes were identified. For this analysis, a percentile rank ≤0.4%, a prediction score ≥0.8, the full IEDB reference set of HLA-class-II alleles, and peptide length of 15 amino acids were applied [105–107]. The shortlisted HTL epitopes were then examined for their ability to induce IL-10, IL-4, and IFN-γ responses employing IL10Pred, IL4Pred, and IFNepitope tools, respectively [108–110]. Both CTL and HTL epitopes were further screened using VaxiJen v2.0 for antigenicity with scores ≥0.4 [102], AlgPred 2.0 for allergenicity with a threshold of <0.4 [111], IEDB Epitope Conservancy Analysis tool for conservancy with a cutoff of ≥90% identity [112], and ToxinPred 2.0 for toxicity with a threshold of <0.6 [113].

Employing the BepiPred 2.0 tool from the IEDB server, the linear B-cell (LBC) epitopes were identified, keeping the cutoff value ≥0.5 and peptide length ≥8 amino acids [114]. To ensure that the selected epitopes were accessible to antibodies, their transmembrane topology was assessed using TMHMM v2.0, and their capacity to induce IgG, IgM, and IgA responses was evaluated using the IgPred server [115,116]. These candidate LBC epitopes were then screened using VaxiJen v2.0 with a threshold of ≥0.4, AlgPred 2.0 with a threshold of <0.4, IEDB Epitope Conservancy Analysis tool with a threshold of ≥90% identity, and ToxinPred 2.0 with a threshold of <0.6 for antigenicity, conservancy, allergenicity, and toxicity, respectively [102,111–113].

### Construction, assessment, and structural refinement of the MEV

The HTL, CTL, and LBC epitopes that were predicted to be antigenic, conserved, having no allergenicity and toxicity, were selected for the MEV construction. Using class-specific GPGPG, KK, and AAY linkers between HTL, LBC, and CTL epitopes, respectively, these epitopes were combined into a single MEV construct [40,45,47,117]. Moreover, using an EAAAK linker, the adjuvant Human β-defensin-3 was fused to the N terminus to induce a better immune response, and a hexahistidine, or a 6×His-tag, was appended to the C terminus of the MEV construct to facilitate a better purification process [118,119].

The overall physicochemical attributes of the final MEV were determined using the ExPASy ProtParam server [120]. Its toxicity, allergenicity, and antigenicity were further evaluated using ToxinPred 2.0 with a threshold of <0.6, AlgPred 2.0 with a threshold of <0.4, and VaxiJen v2.0 with a cutoff score of ≥0.4, respectively [102,111,113]. The MEV secondary-structure was predicted using PSIPRED 4.0 [121,122]. The tertiary-structure of the MEV was modelled using I-TASSER, and the model with the highest estimated TM-score among the five generated structures was selected for subsequent structural refinement [123]. The selected model was refined using GalaxyRefine through the GalaxyWEB server to improve backbone geometry and side-chain packing [124]. The refined structure was finally assessed using the SWISS-MODEL structure validation, PROCHECK, and ProSA-web [92,93,95,96].

To improve structural stability, disulfide engineering was carried out using Disulfide by Design 2 (DbD2, version 2.12) [125]. Candidate residue pairs that had disulfide bond energies below 2.5 kcal/mol were retained. Additional filters, including a χ3 dihedral angle of –87° or +97° (with an ±30° allowed deviation) and a 114.6° ± 10° Cα-Cβ-Sγ bond angle, were applied, reflecting the most commonly observed geometries of native disulfide bonds [117,125,126]. Afterwards, using the ElliPro server, keeping ≥0.5 as the cutoff score, conformational B-cell epitopes from the MEV structure were identified, based on the principle that antibody recognition generally favors solvent-exposed surface residues rather than buried regions [127,128].

### MEV-TLR4 interaction analysis, immune simulation, and *in silico* cloning

To examine whether the host immune system engages the designed MEV, protein-protein docking was performed between the MEV and human TLR4 using ClusPro 2.0 server [129]. The server uses rigid-body docking by sampling multiple relative orientations of the two proteins and clustering low-energy poses based on pairwise RMSD [129]. The TLR4 structure (PDB ID: 4G8A) was retrieved from the RCSB Protein Data Bank and prepared in BIOVIA Discovery Studio with standard optimization procedures [129]. Using the PDBsum server protein-protein interaction tool, intermolecular contacts including hydrogen bonds, salt bridges, and interface residues within the selected MEV-TLR4 complex were analyzed [130]. The final docked complex and its interacting residues were then visualized in PyMOL v3.1 [131].

Binding affinity of the docked MEV-TLR4 complex, including dissociation constant (Kd) and binding free energy (ΔG), were estimated using the PRODIGY server [132].

Furthermore, the flexibility, stability, and collective motions of the complex were examined utilizing the iMODS server, which provides normal mode and molecular dynamics-based assessments of macromolecular flexibility [133].

To estimate the immunological potential of the constructed vaccine, immune simulation was performed *in silico* using the C-ImmSim server with three vaccine doses of 1,000 antigen units without lipopolysaccharides, administered at 1, 84, and 168 time-steps [134]. The simulation was run with 1,050 total time-steps, a system volume of 10 µL, and a random seed of 12345 [119,134].

The amino acid sequence of the designed MEV was then reverse translated and codon-optimized for *Escherichia coli* strain K-12 using the Java Codon Adaptation Tool (JCAT) [135]. During optimization, codon usage was adjusted to match host preference while avoiding undesirable sequence features such as internal ribosome-binding sites, transcription terminators, and inappropriate restriction enzyme recognition sites. To evaluate its suitability for bacterial expression, we calculated the GC content of the optimized sequence and its codon adaptation index (CAI) keeping target values of GC content between 30% and 70% and CAI > 0.8 [135,136]. With appropriate restriction sites introduced for insertion into the *E. coli* expression vector pET-28a (+) backbone, the optimized MEV sequence was cloned *in silico* using SnapGene v8.2.2 [137–139].

## Results

Using an integrative subtractive genomics and reverse vaccinology strategy, we screened the complete proteome of *P. multocida* Pm70 to identify proteins with potential value as both drug and vaccine targets. The complete filtering progression is briefly described in **Table 1**.

**Table 1.**
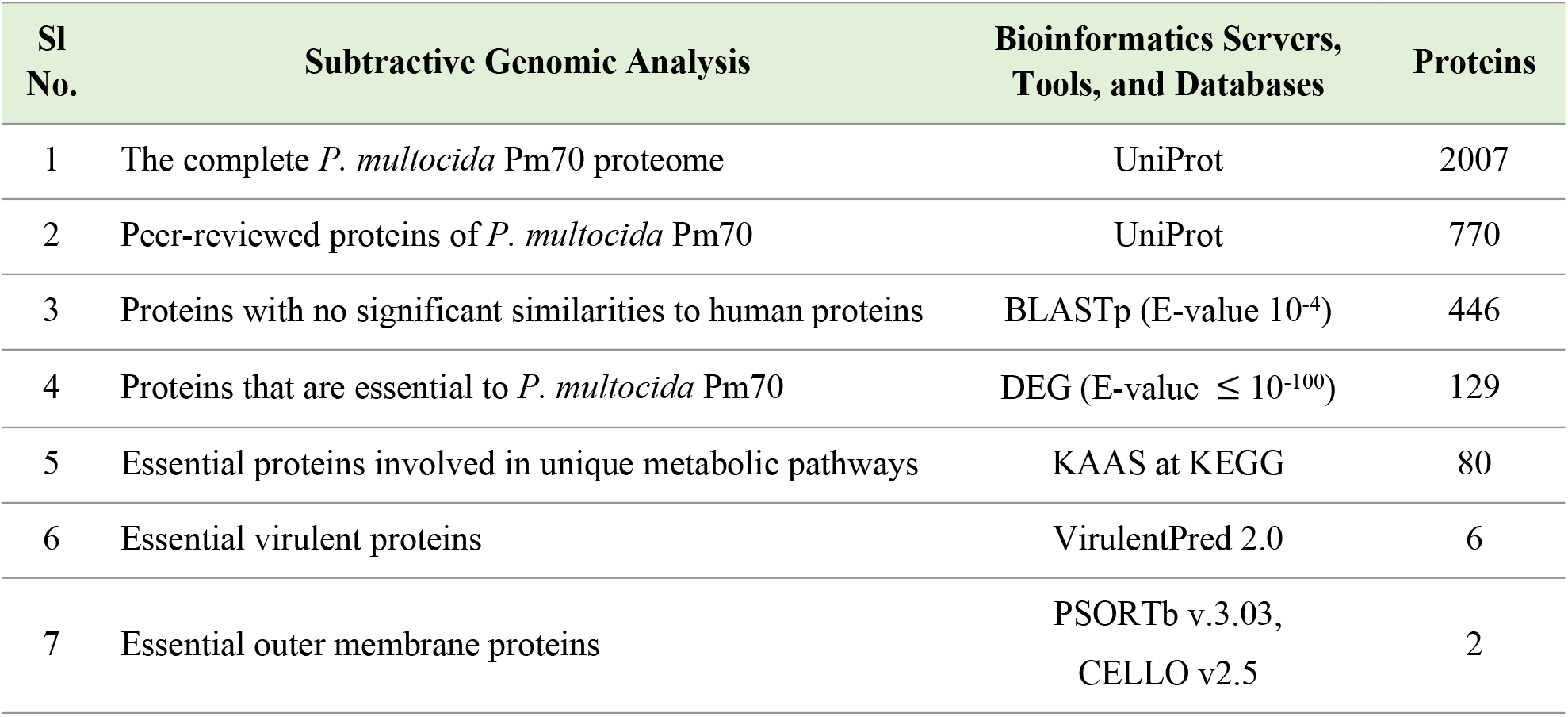
The subtractive genomic analysis of *P. multocida* Pm70, showing sequential filtering of the entire proteome with the corresponding bioinformatics servers, tools, and databases, along with the total proteins retained at each step.

### Retrieval of protein sequences and screening of non-homologous and essential proteins

To effectively minimize overrepresentation of certain proteins, reduce redundancy in the proteome, and ensure the representativeness and accuracy of the data, a total of 770 peer-reviewed proteins from the complete *P. multocida* Pm70 proteome, comprising 2,007 entries in total, were retrieved and subjected to stepwise subtractive genomic screening. Comparison against the human proteome retained 446 (57.92%) proteins with no significant similarities with human proteins, excluding the rest. Subsequent analysis of the retained non-homologous sequences identified 129 proteins that are indispensable for the pathogen’s survival (**S1 Table**). The complete filtering progression is briefly described in **Table 1**.

### Exploration of metabolic pathways, known drug target investigation, virulence association, and subcellular localization

The essential proteins were subsequently examined for their association in important metabolic pathways. Manual comparison between the human and pathogen pathways identified 3 proteins involved in a single pathway common to both, which were consequently excluded from further analysis. Of the remaining targets, 80 proteins were found to be associated with 60 pathogen-specific pathways, while the remaining 46 proteins were pathway-independent but carried out important biological functions. The unique pathways and the number of associated proteins is summarized in **Table 2**.

**Table 2.**
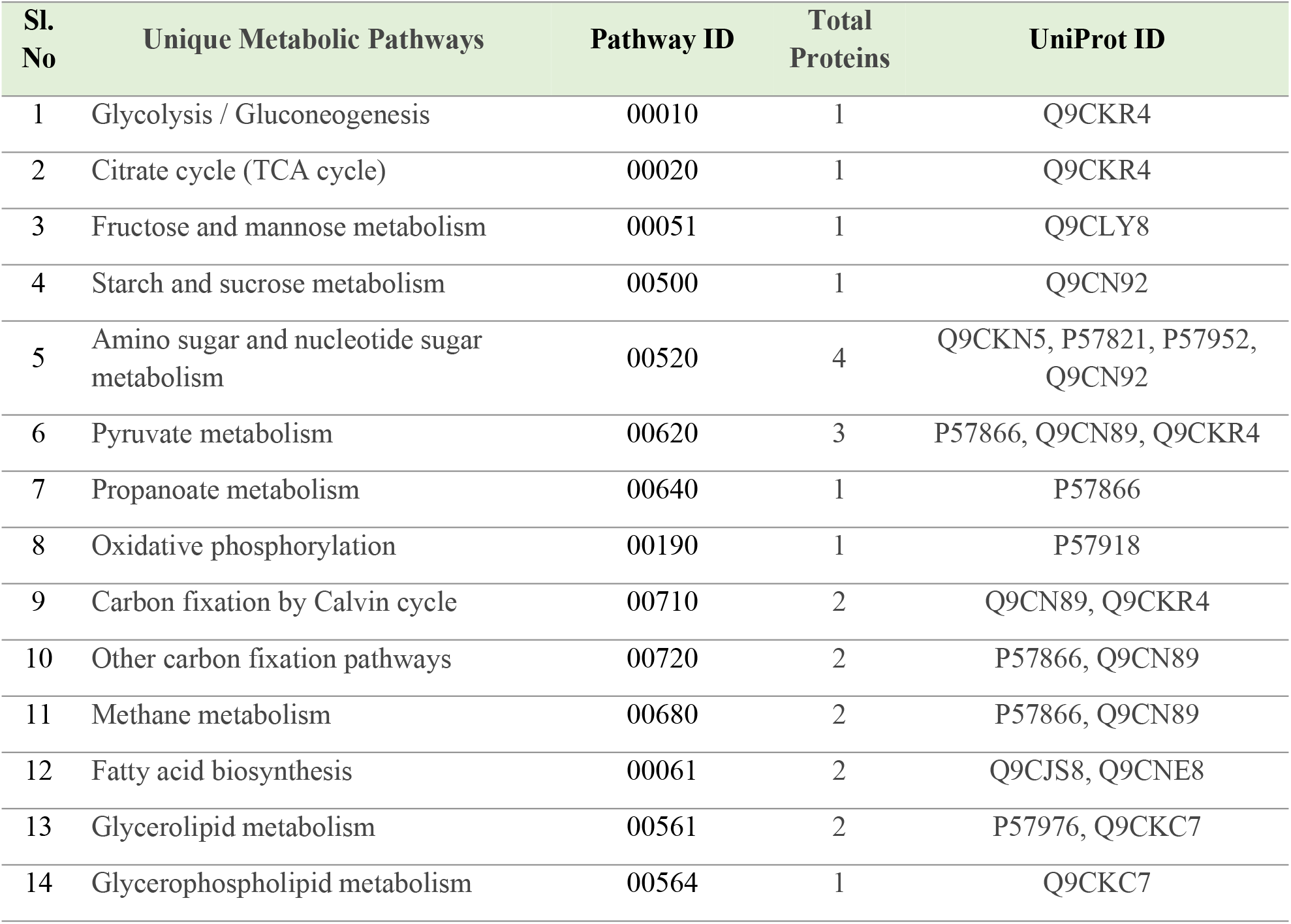
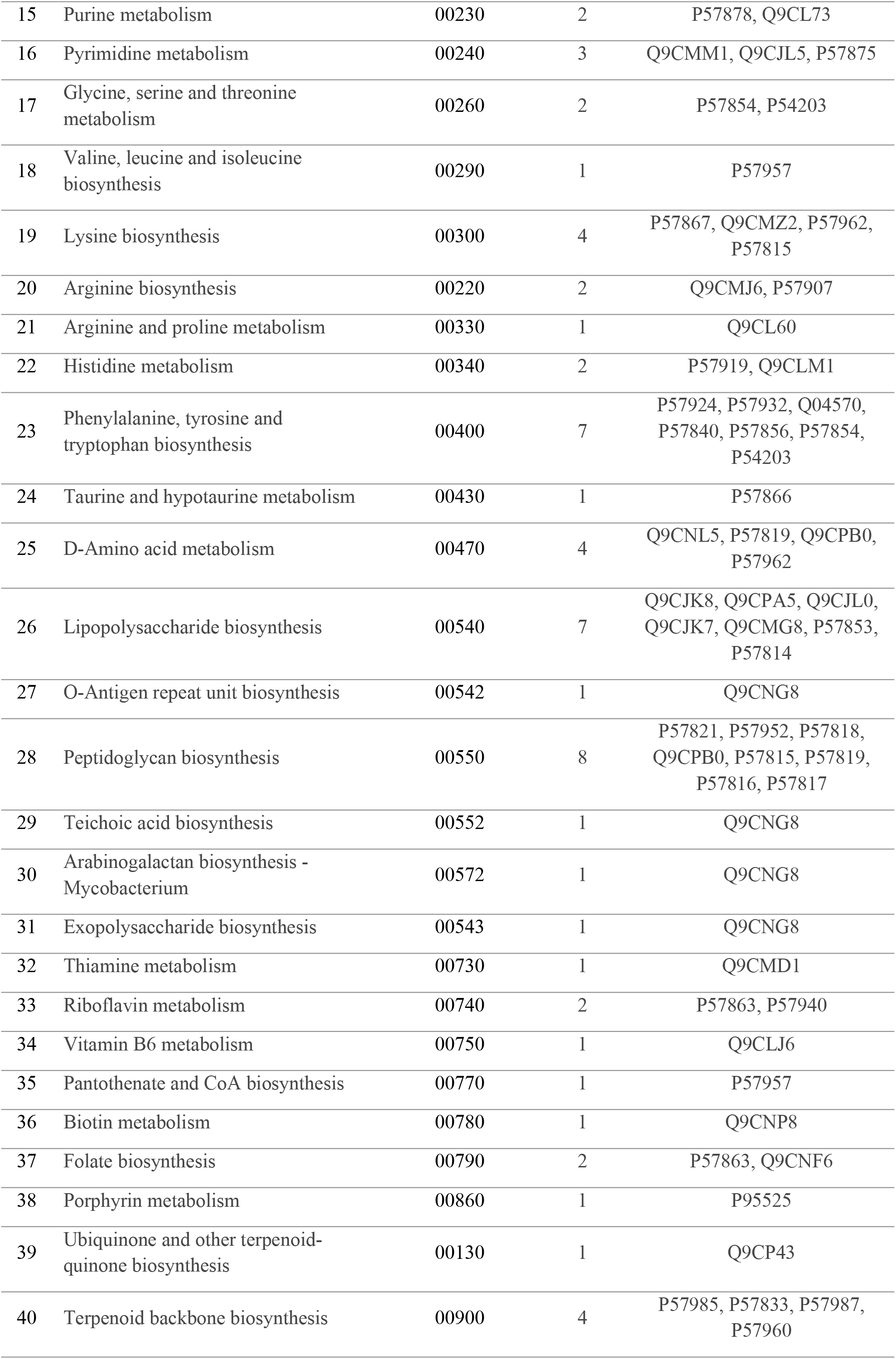
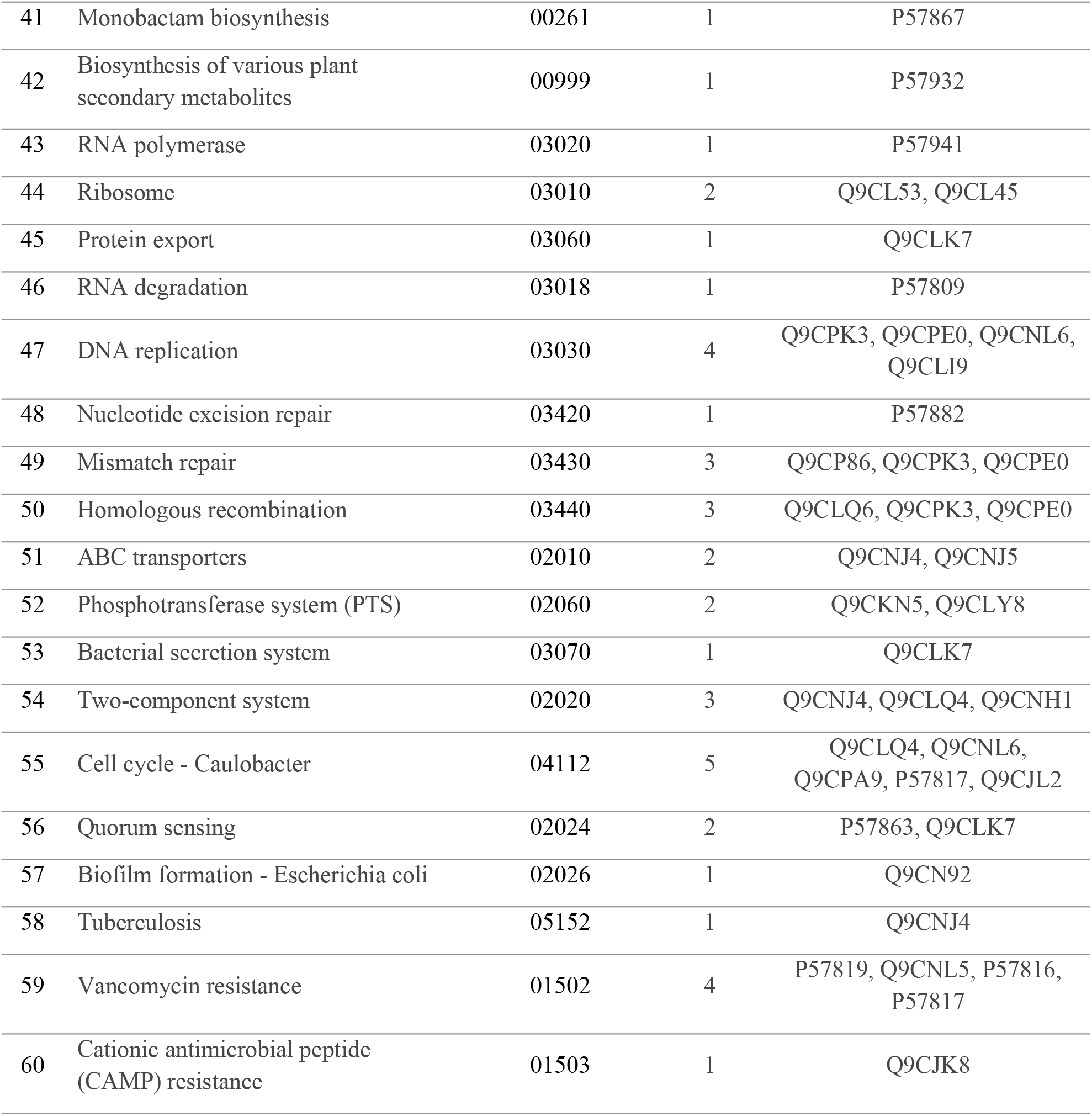
Metabolic pathways unique to *P. multocida*, listing KEGG pathway IDs, the number of proteins involved in each pathway and their corresponding UniProt accessions.

Homology screening against known drug targets identified 49 proteins homologous to entries deposited in the DrugBank database (version 6.0). The remaining 77 proteins were classified as novel targets due to their lack of similarity with existing entries. Among these, 35 novel drug targets were involved in 44 pathogen-specific pathways, including lipopolysaccharide biosynthesis, two-component system, D-amino acid metabolism, quorum sensing, vancomycin resistance, bacterial secretion system, biofilm formation, and peptidoglycan biosynthesis (**S2 Table**). In addition, 42 proteins from the pool of 46 pathway-independent proteins were identified as novel drug targets that govern critical biological functions, such as cell division, replication, translation, transcription, repair, drug efflux, and host colonization (**S3 Table**).

Virulence prediction further identified six essential proteins associated with virulence-related functions in *P. multocida* (**S4 Table**). Subsequent subcellular localization analysis classified the essential proteins across the cytoplasm, periplasm, inner-membrane, extracellular, and outer-membrane compartments, as illustrated in **Fig 2** (**S5 Table**). The proteins displaying outer membrane characteristics predicted by both tools are listed in **Table 3**. Two proteins, OMP47 and LptD, emerged as compelling candidates, simultaneously satisfying all major prioritization criteria as they lacked homology to human proteins, were essential to the pathogen’s viability, and were localized to the bacterial outer membrane.

**Fig 2.**
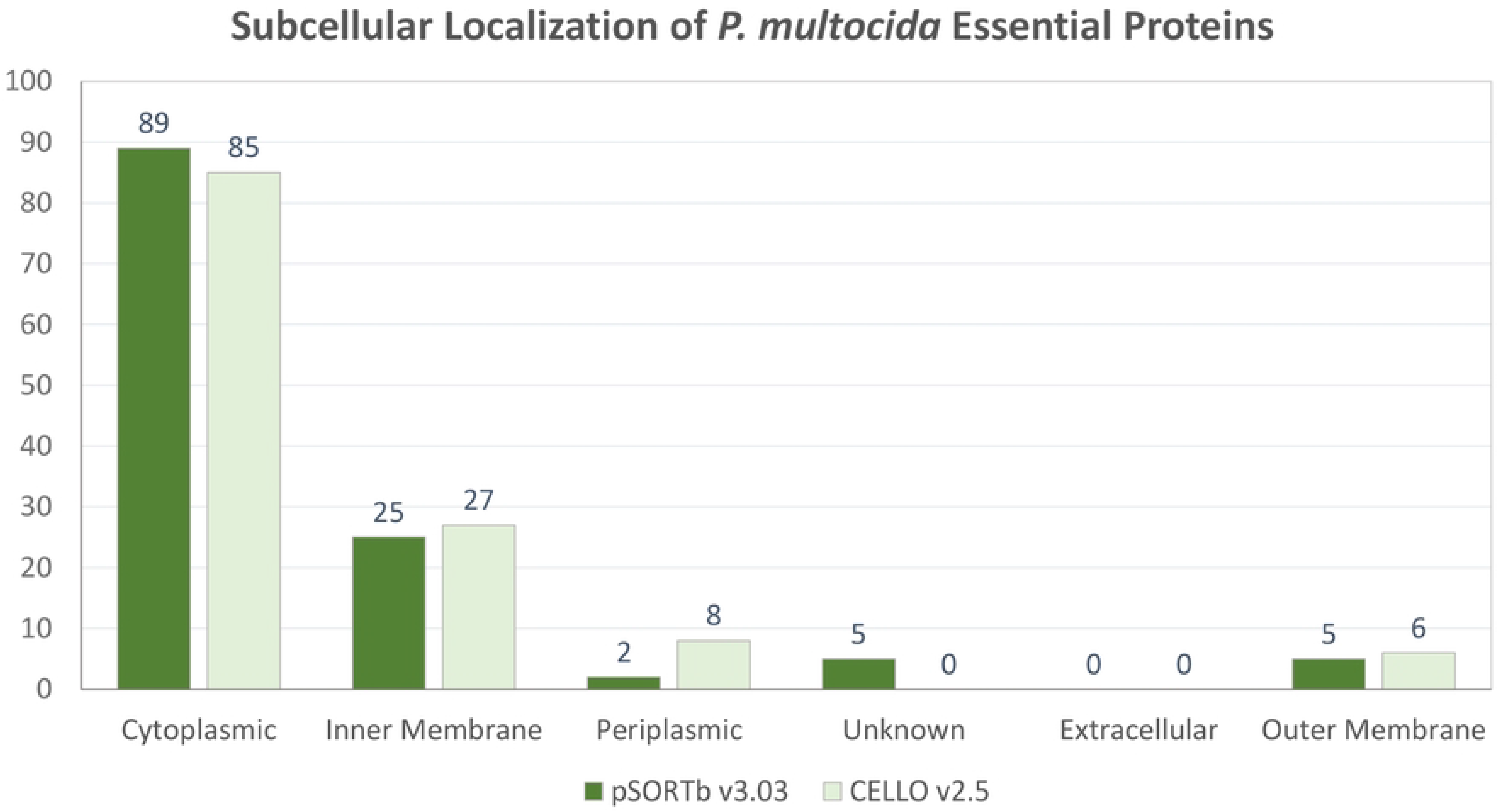
The subcellular localization of the essential proteins. The black and grey bars climbing upwards along the y-axis indicate the number of proteins predicted by pSORTbv3.03 and CELLO v2.5, respectively. At the same time, the x-axis holds the categories of different subcellular localizations.

**Table 3.**
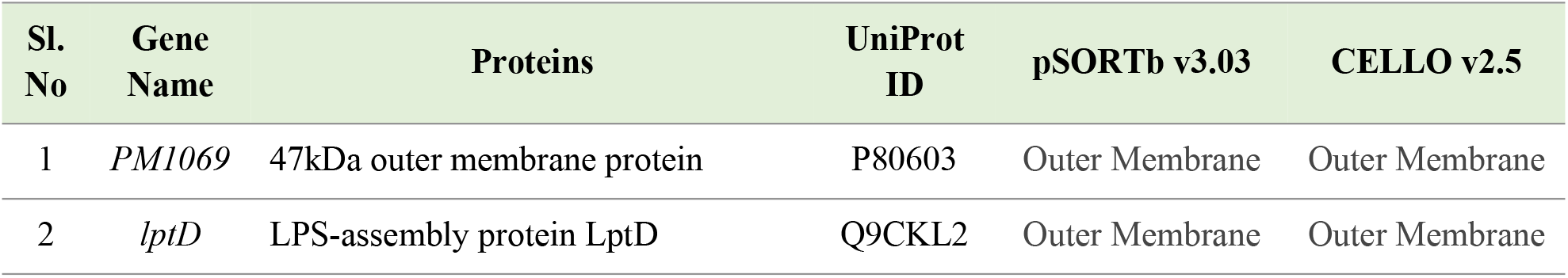
Outer membrane proteins predicted by both pSORTbv3.03 and CELLOv2.5 subcellular localization tools.

### Physicochemical characterization, protein interaction analysis, and structural modelling

The physiochemical property analysis revealed distinct structural attributes for both outer membrane proteins, as detailed in **Table 4**. OMP47 and LptD comprise 428 and 782 amino acids, with calculated molecular weights of 46,061.23 Da and 90,647.20 Da, respectively. Both candidates are highly basic in nature, as indicated by their theoretical isoelectric points of 9.12 (OMP47) and 9.03 (LptD). Negative grand average of hydropathicity (GRAVY) values (–0.100 for OMP47 and –0.708 for LptD) confirmed their overall hydrophilic character, while high aliphatic indices (82.87 and 73.63, respectively) indicated strong thermal stability. The functional connectivity profiles according to the STRING network analysis revealed statistically significant interaction networks for both targets, as shown in **Fig 3**, indicating PPI enrichment *p*-values of 0.0274 for OMP47 and <1.0 x 10^-16^ for LptD. OMP47 established a network containing 7 nodes and 12 edges, functionally interacting with six core proteins BamA, PM0336 (HgbB), PM1070, PM1066, and MutH, which are involved in outer membrane related functions within *P. multocida*, alongside Dat1, which is involved in cellular defense. In contrast, LptD demonstrated a highly complex network spanning 21 nodes and 93 edges, interacting with 20 neighbouring proteins. These primarily included LptE, LptA, and LptC which are integral to lipopolysaccharide translocation and assembly, alongside BamA, BamD, BamE, BamC, TolR, TolB, FtsQ, and FtsL which are essential for outer membrane integrity and cell division. The STRING network statistics are listed in **Table 4**, and the biological functions of all interacting network proteins are detailed in **S6 Table**.

**Fig 3.**
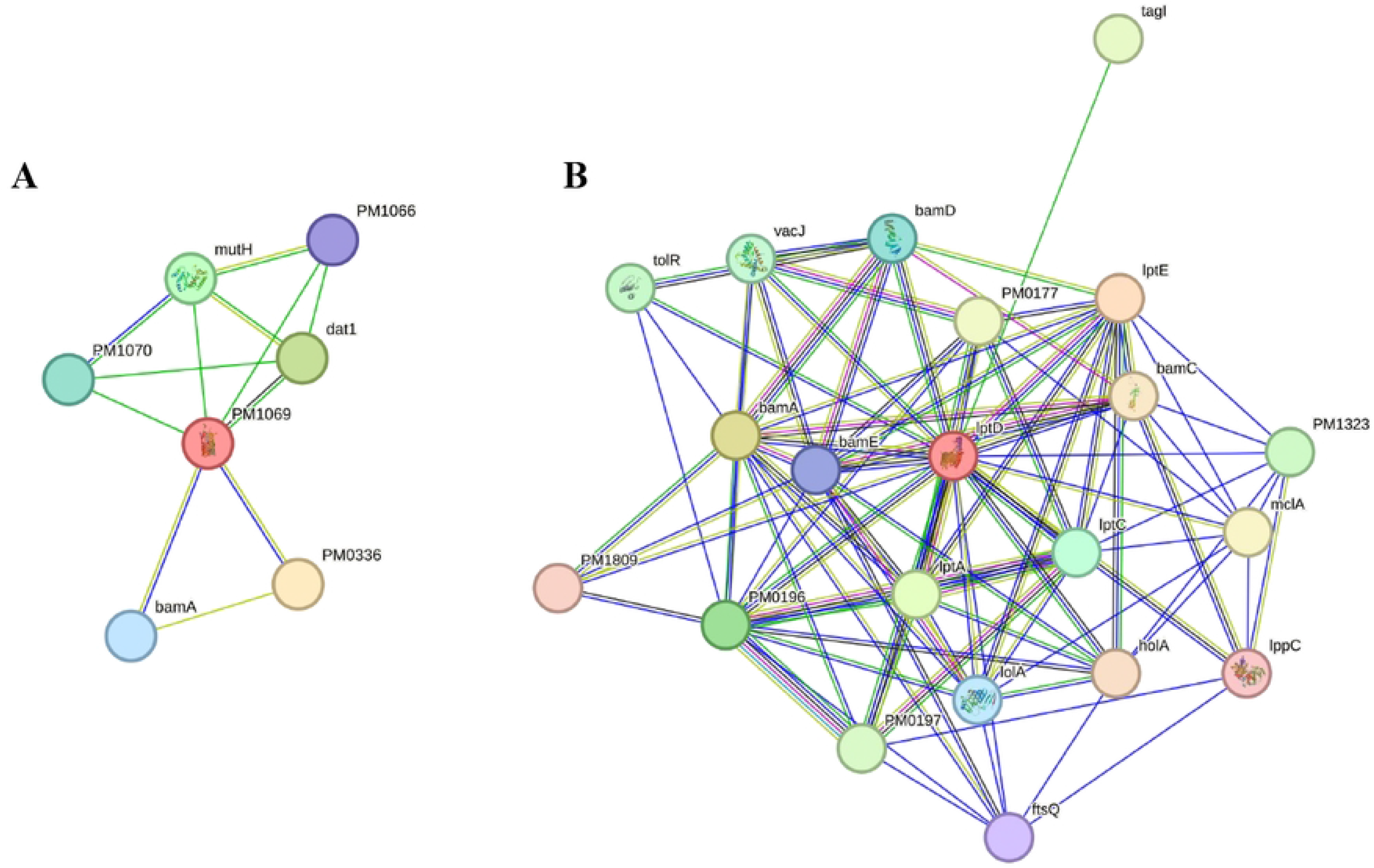
Protein-protein interaction networks of the proteins OMP47 and LptD. The queried proteins (**A**) OMP47 and (**B**) LptD are shown as red nodes, and each node represents a protein encoded by a single gene locus. Filled nodes indicate proteins with known or predicted three-dimensional structures, whereas unfilled nodes indicate proteins with no available structural information. Edges represent protein-protein associations.

**Table 4.**
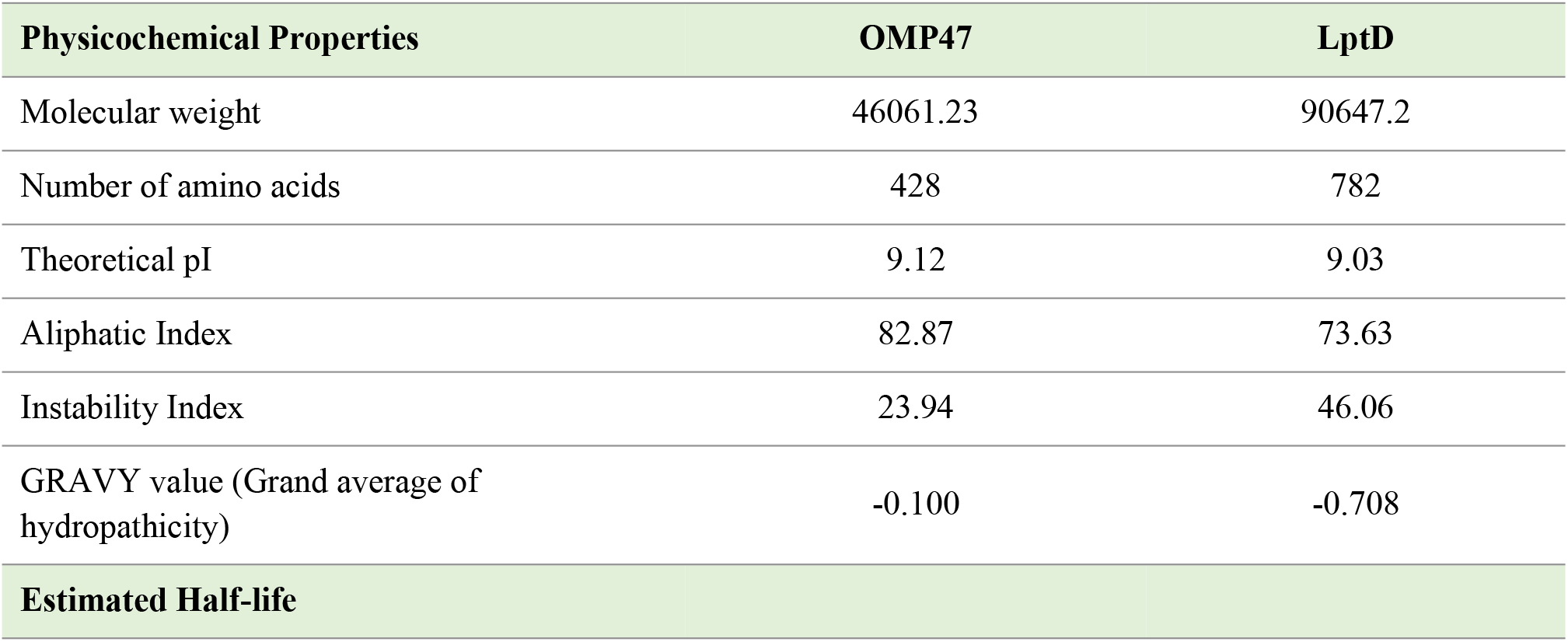
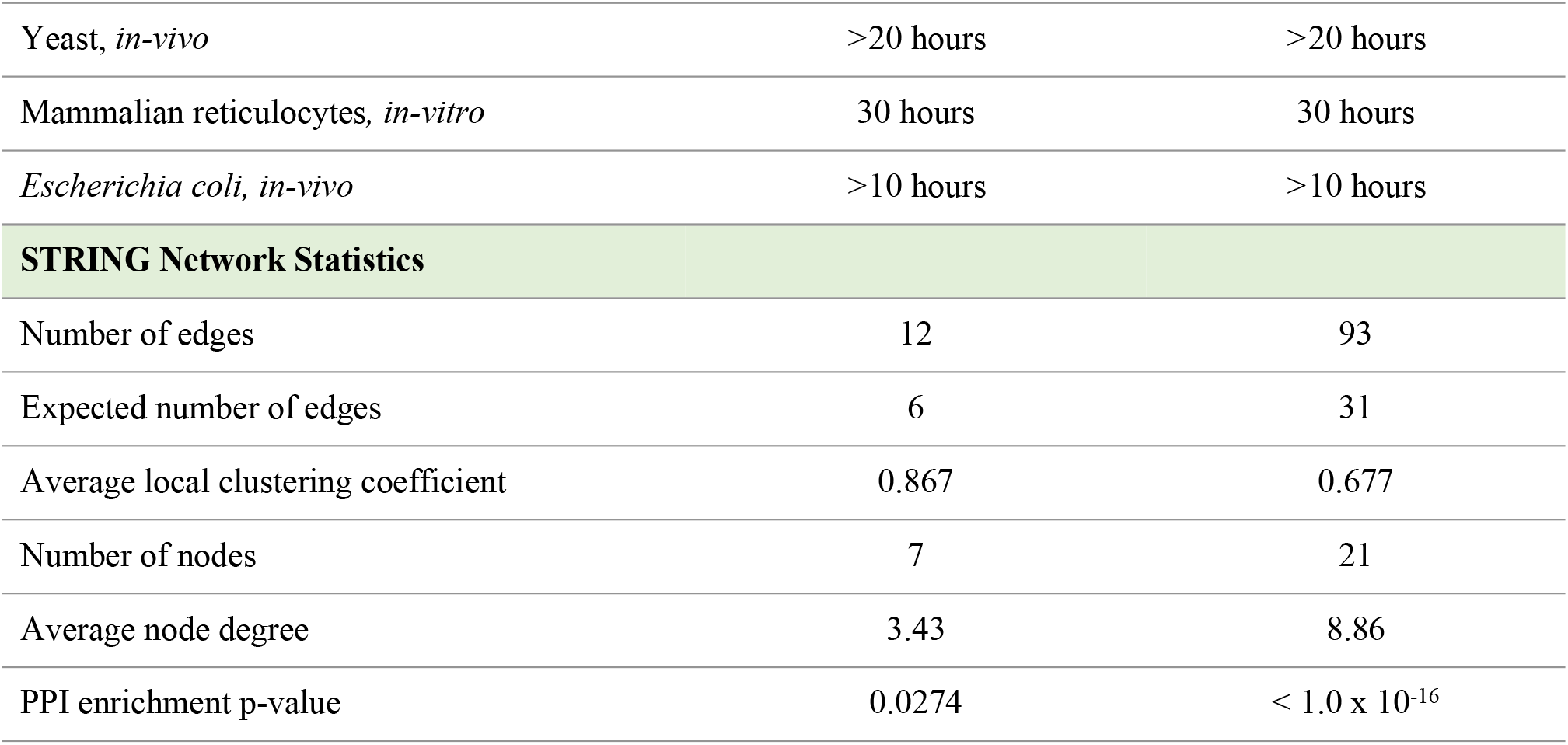
Physiochemical properties and STRING protein-protein interaction network statistics of the outer membrane proteins, OMP47 and LptD.

The three-dimensional models of the two outer membrane proteins, OMP47 and LptD, were generated using SWISS-MODEL server (illustrated in **Fig 4** and **Fig 5**, respectively) and cross-validated against the AlphaFold Protein Structure Database to ensure high structural consistency. Structural assessment of the SWISS-MODEL generated three-dimensional models showed that 96.24% of OMP47 residues and 96.15% of LptD were located in the favored region of the Ramachandran plot. This high quality was further supported by low MolProbity scores (1.05 for OMP47 and 0.79 for LptD) and high ERRAT quality factors (94.35 for OMP47 and 92.62 for LptD). PROCHECK analysis further supported the quality of the three-dimensional models, with 91.1% of OMP47 residues and 91.1% of LptD in the favored regions. In addition, ProSA-web yielded Z-scores of −6.64 for OMP47 and −4.6 for LptD (**S1 Fig**), confirming that both structures fall well within the typical energy range of experimentally resolved native proteins of similar size.

**Fig 4.**
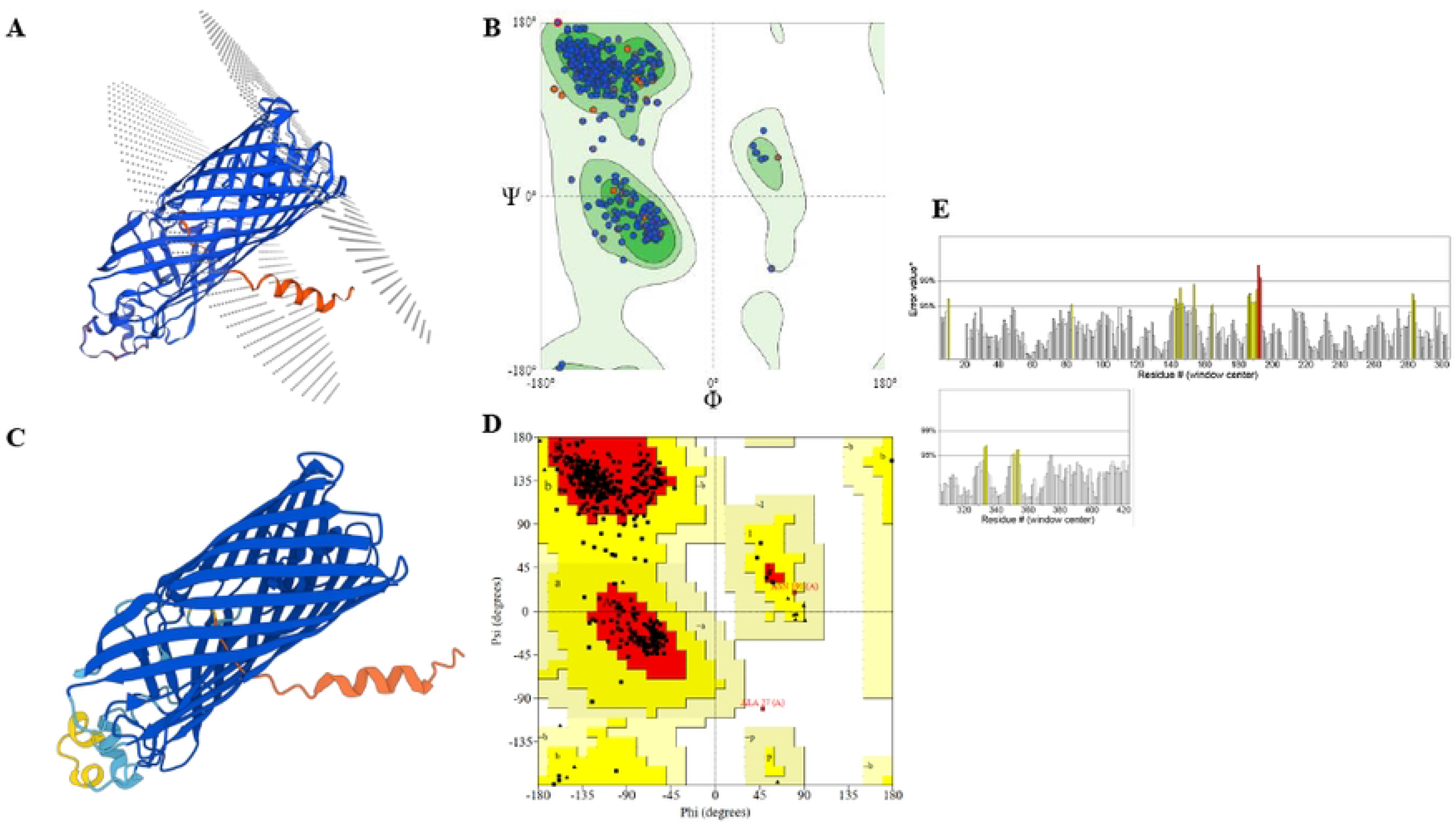
Structural modelling and validation of OMP47. (**A**) SWISS-MODEL generated three-dimensional structure of OMP47. (**B**) SWISS-MODEL Structure Evaluation Ramachandran plot of the SWISS-MODEL generated three-dimensional structure of OMP47. (**C**) AlphaFold three-dimensional structure of OMP47. (**D**) PROCHECK Ramachandran plot of the SWISS-MODEL generated three-dimensional structure of OMP47. (**E**) ERRAT residue-wise error profile of the generated three-dimensional structure of OMP47.

**Fig 5.**
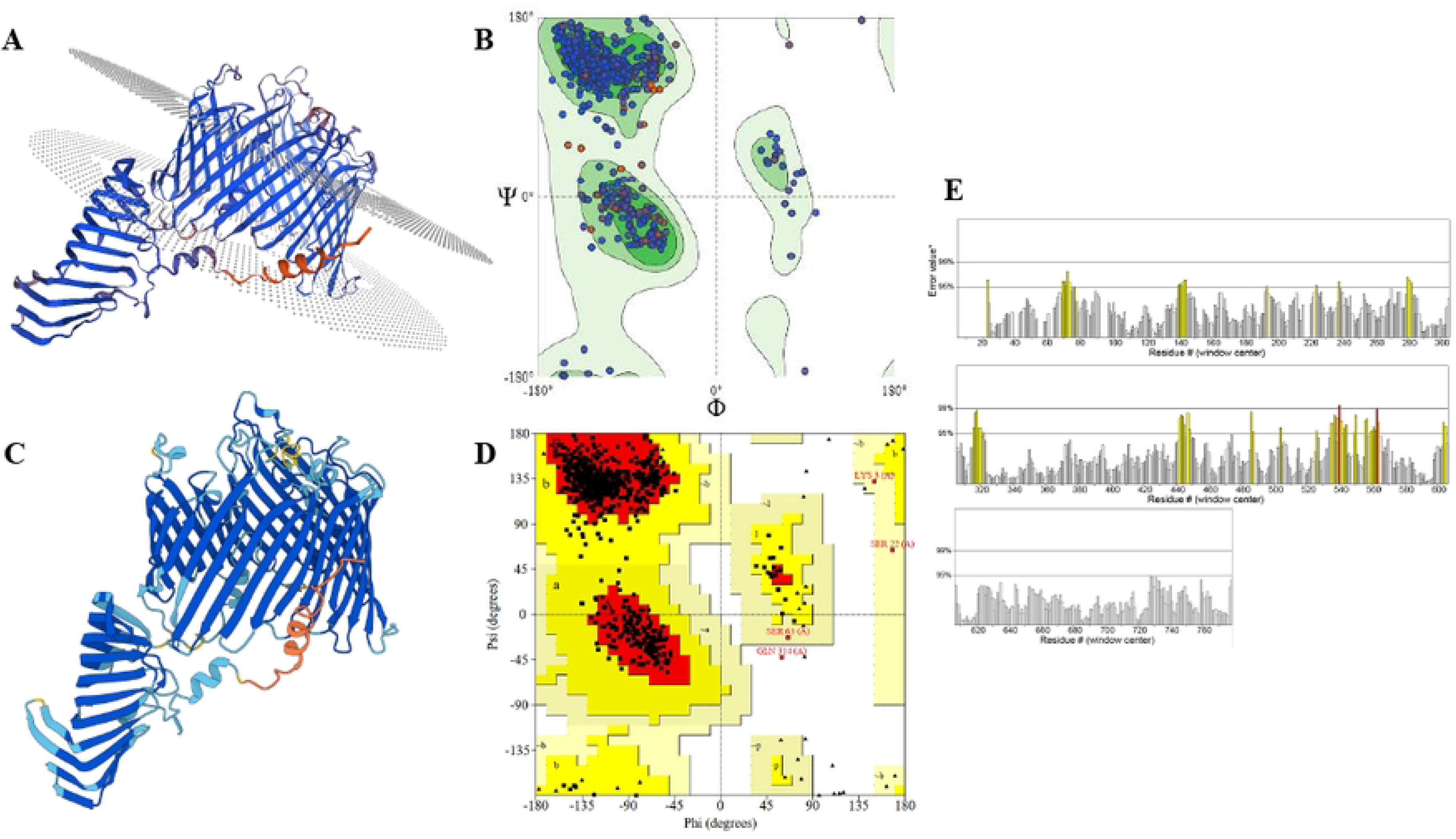
Structural modelling and validation of LptD. (**A**) SWISS-MODEL generated three-dimensional structure of LptD. (**B**) SWISS-MODEL Structure Evaluation Ramachandran plot of the SWISS-MODEL generated three-dimensional structure of LptD. (**C**) AlphaFold three-dimensional structure of LptD. (**D**) PROCHECK Ramachandran plot of the SWISS-MODEL generated three-dimensional structure of LptD. (**E**) ERRAT residue-wise error profile of the SWISS-MODEL generated three-dimensional structure of LptD.

### Binding-site prediction, ligand identification, and molecular docking

Binding-site analysis identified 10 potential binding pockets in OMP47 and 35 in LptD, as depicted in **Fig 6**. Binding pockets were prioritized based on drug score, volume, depth, and residue organization (**S7 Table**), with the top-ranked OMP47 pocket comprising 53 residues, and the LptD pocket comprising 70 residues (**S8 Table**).

**Fig 6.**
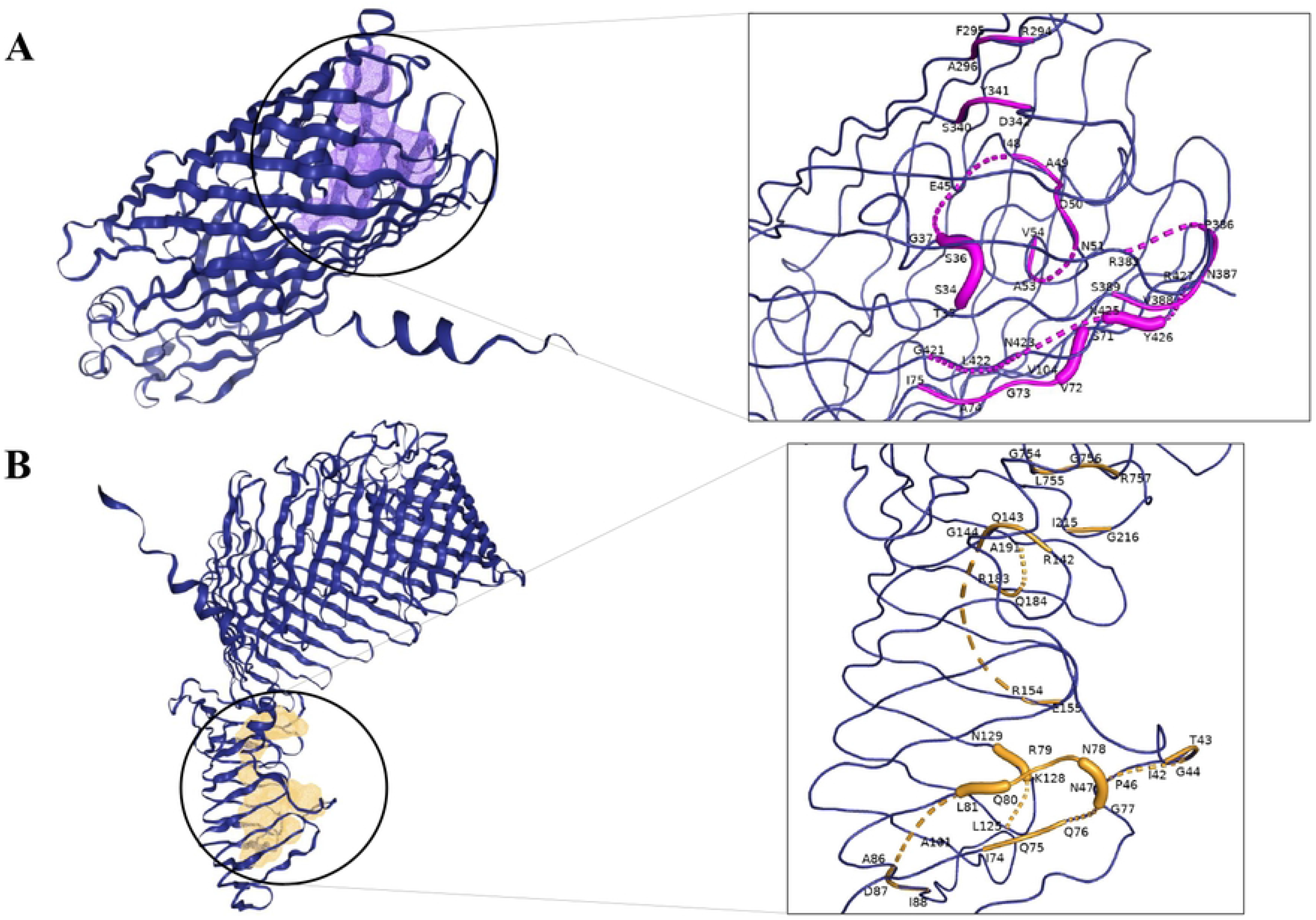
Predicted ligand-binding pockets in OMP47 and LptD. (**A**) OMP47 showing the highest-scoring binding pocket (left, shaded region) and an enlarged view of the cavity with lining residues labelled (right, magenta). (**B**) LptD with the top-ranked binding pocket highlighted (left, shaded region) and a zoomed view depicting the pocket-lining residues (right, orange).

Based on pocket architecture and compatibility, alpha-D-galactopyranuronic acid (ADA) and (hydroxyethyloxy)tri(ethyloxy)octane (C8E) were selected as representative docking ligands for OMP47 and LptD, respectively (**S2 Fig**).

Molecular docking predicted binding energies of –6.1 kcal/mol for the OMP47-ADA complex and –5.3 kcal/mol for the LptD-C8E complex, suggesting stronger binding of ADA to OMP47. As illustrated by the two-dimensional and three-dimensional interaction profiles in **Fig 7**, post-docking analysis revealed that OMP47 formed hydrogen bonds with ADA through residues Arg350, Thr348, Arg383, Asp342, and Tyr288. Conversely, LptD formed hydrogen bonds with C8E through residues Asp111, Arg79, Tyr109, Lys128, Gln76, and Thr130. In both complexes, the ligands occupied the predicted binding pockets and established stabilizing interactions with key surrounding residues.

**Fig 7.**
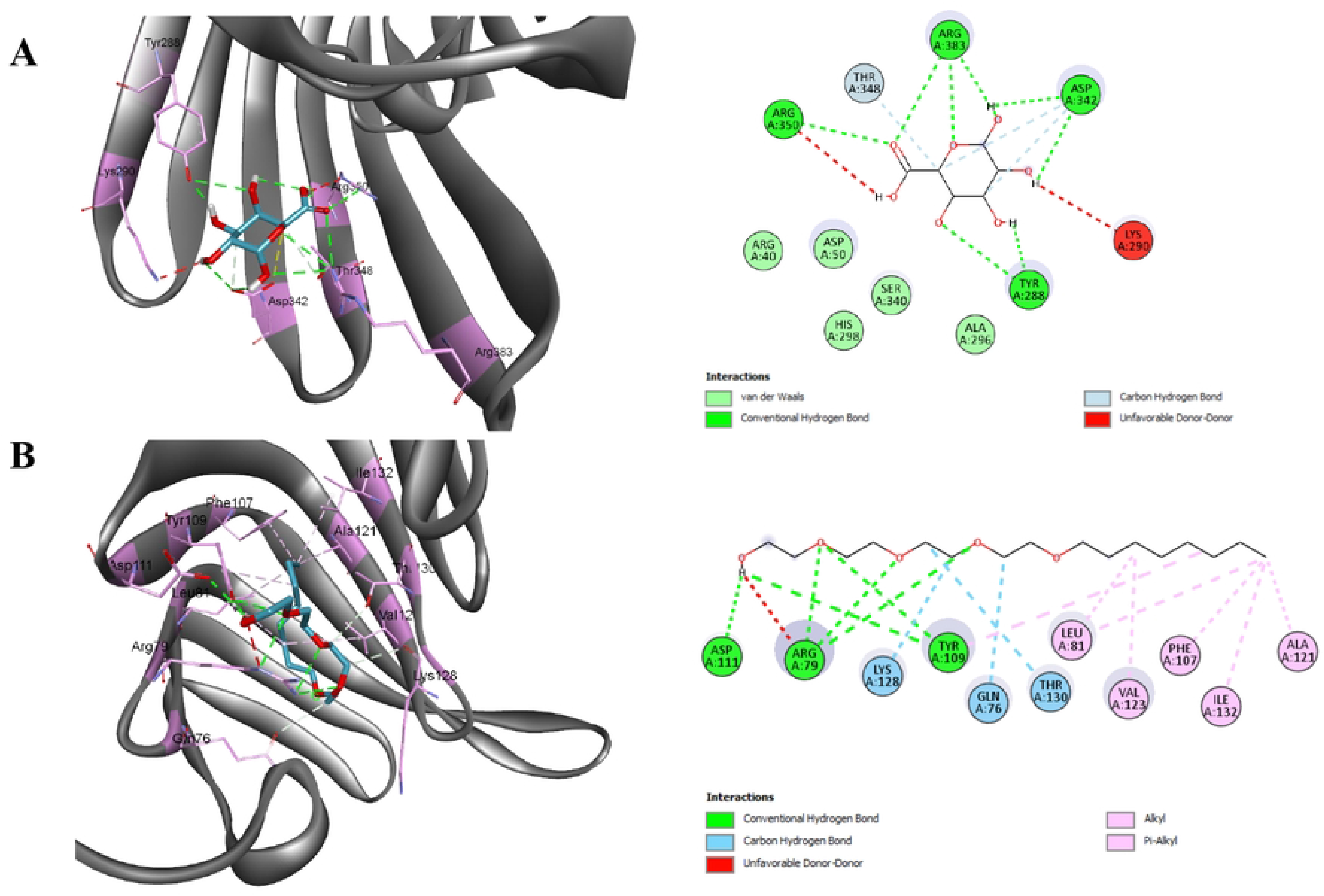
Molecular docking analysis of the selected ligands with OMP47 and LptD. (**A**) The OMP47-ADA complex two-dimensional and three-dimensional interaction profiles. (**B**) The LptD-C8E complex two-dimensional and three-dimensional interaction profiles. In the three-dimensional panels, the ligand-binding residues are shown in stick representation together with the bound ligands. In the two-dimensional panels, dotted lines indicate the predicted interactions formed within each protein-ligand complex.

### Antigenicity prediction and epitope mapping

Because OMP47 and LptD are outer membrane-associated and showed favorable VaxiJen antigenicity scores of 0.6614 and 0.6820, respectively, both proteins were selected as target antigens for epitope mapping. CTL epitope prediction initially yielded 22,653 candidate peptides from OMP47 and 41,769 from LptD. Progressive prioritization reduced these to 19 antigenic, conserved, non-allergenic, non-toxic, and immunogenic CTL candidates (4 from OMP47 and 15 from LptD). From this prioritized panel, seven CTL epitopes (1 from OMP47 and 6 from LptD) were finally selected for inclusion in the MEV construct, as detailed in **Table 5**.

**Table 5.**
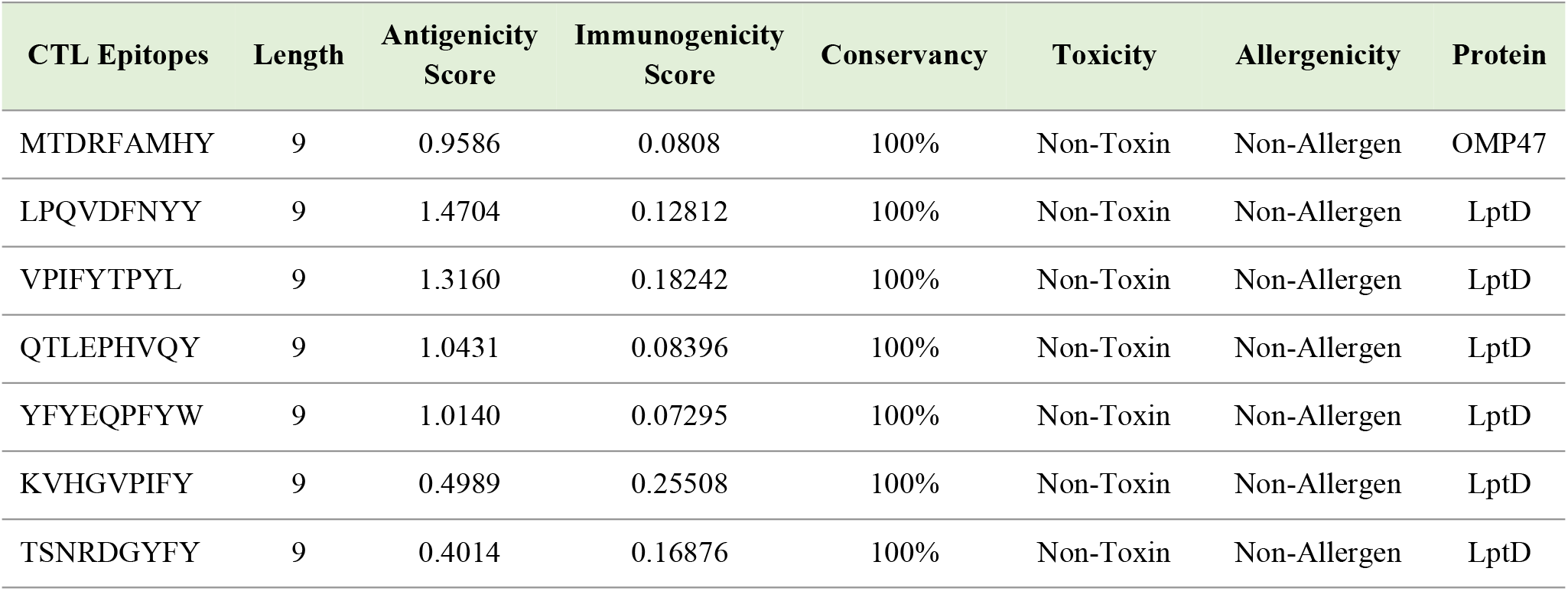
Prioritized CTL epitopes predicted from OMP47 and LptD for construction of the MEV with immunogenicity scores, antigenicity values, predicted toxicity and allergenicity profiles.

HTL epitope prediction identified 11,178 candidates from OMP47 and 20,736 from LptD. After applying stringent threshold filters for antigenicity, conservancy, allergenicity, and toxicity, 47 high-confidence HTL epitopes were retained (14 from OMP47 and 33 from LptD). Further characterization of their specific cytokine-inducing profiles narrowed this selection down to 12 prioritized HTL epitopes (5 from OMP47 and 7 from LptD) for vaccine construction, as presented in **Table 6**.

**Table 6.**
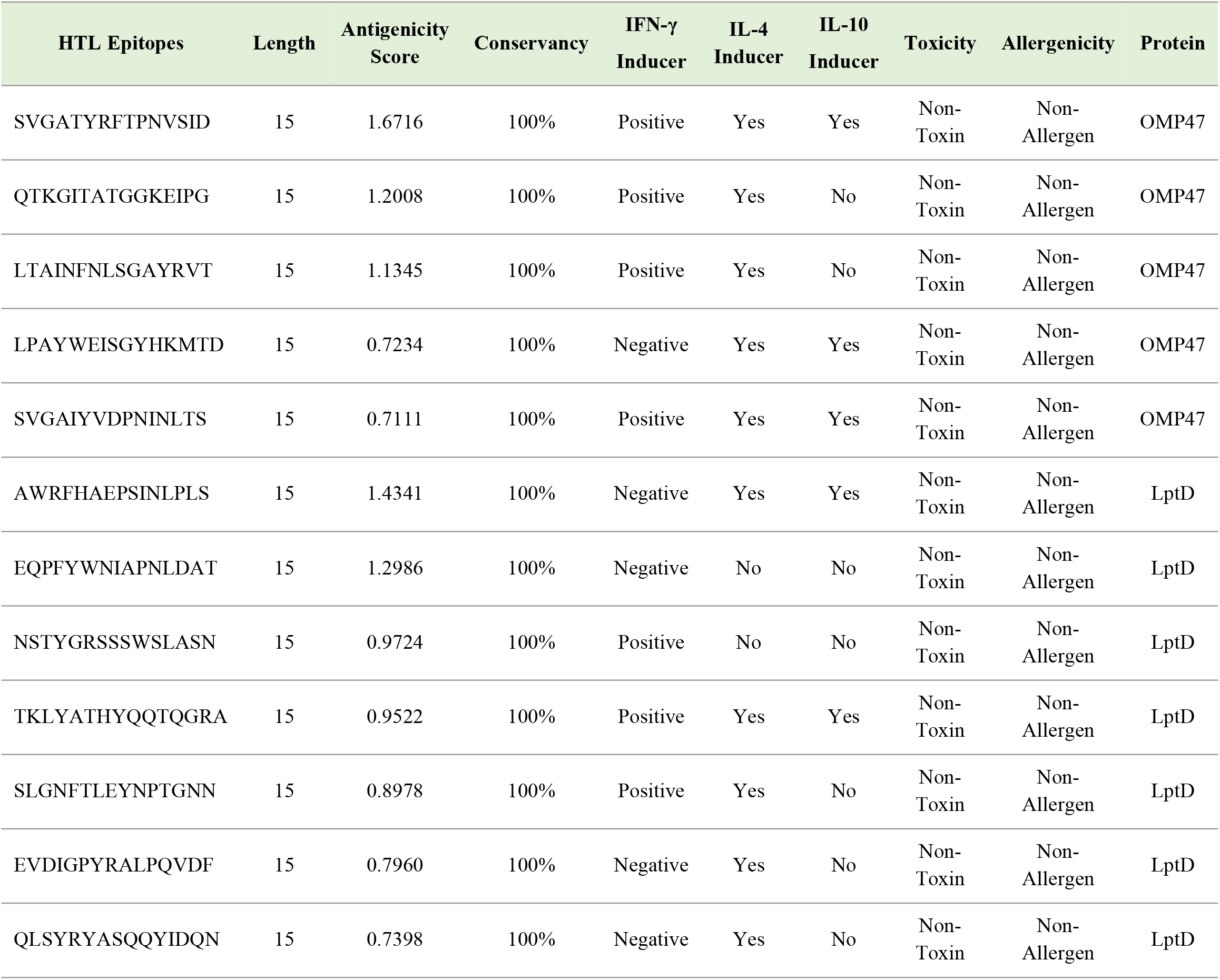
Prioritized HTL epitopes predicted from OMP47 and LptD for construction of the MEV with antigenicity scores, predicted toxicity and allergenicity, and IL-4, IFN-γ, and IL-10 induction profiles.

LBC epitope screening identified 22 candidates, including 7 from OMP47 and 15 from LptD. Among these, seven epitopes met the criteria of antigenicity, conservancy, non-allergenicity, and non-toxicity. Transmembrane topology analysis demonstrated that two OMP47 epitopes and one LptD epitope were exposed on the extracellular surface, making them accessible for antibody engagement, while the remaining sequences were located on the inner side of the membrane. All seven LBC epitopes (3 from OMP47 and 4 from LptD) were retained as candidates for the MEV construct, as shown in **Table 7**. The full dataset of all predicted CTL, HTL, and LBC epitopes are provided in **S11**, **S12**, and **S13 Tables**, respectively.

**Table 7.**
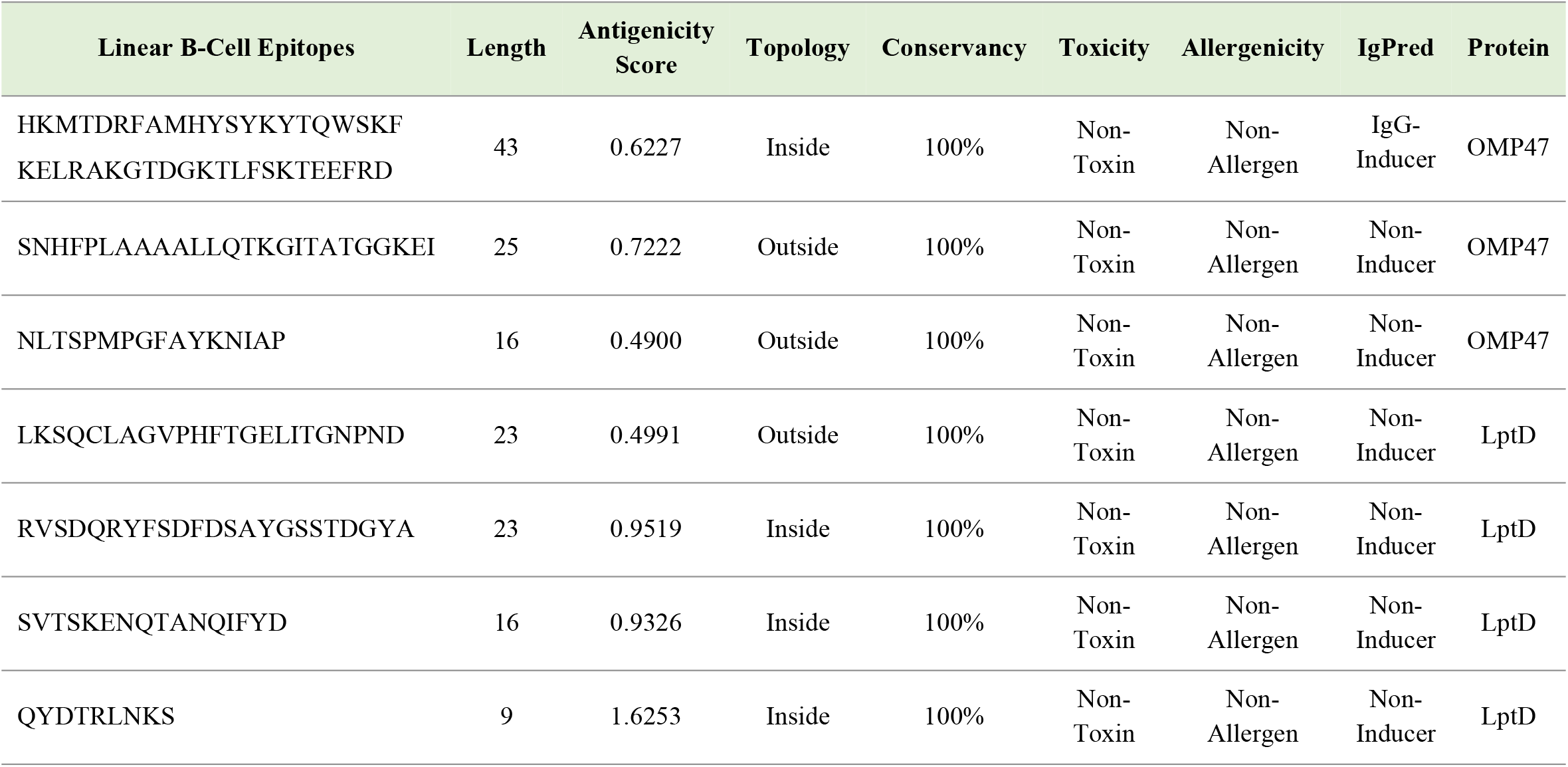
Prioritized LBC epitopes predicted from OMP47 and LptD for construction of the MEV with antigenicity scores, predicted toxicity and allergenicity, transmembrane topology, and immunoglobulin inducing profiles.

### Construction, assessment, and structural refinement of the MEV

The final MEV construct incorporated 26 prioritized epitopes, comprising of 12 HTL, seven LBC, and seven CTL epitopes, which was assembled into a single chimeric sequence using class-specific linkers, an N-terminal adjuvant, and a C-terminal 6×His-tag (**S3 Fig)**. The resulting construct was 552 amino acids in length, with a molecular weight of 60.6kDa, a theoretical pI of 9.39, and an instability index of 34.41, suggesting that it possesses a net positive charge and remains highly stable under physiological conditions. Its aliphatic index and negative GRAVY value further indicated a relatively thermostable and hydrophilic profile, supporting its compatibility with soluble bacterial expression systems. Total charge composition analysis further supported the basic nature of the construct. Furthermore, VaxiJen antigenicity score of 0.9486 indicated the construct to be highly antigenic, while AlgPred2.0 and ToxinPred2 classified it as non-allergenic and non-toxic, respectively (**S9 Table**).

Secondary-structure analysis showed that random coils predominated, with smaller proportions of β-strands and α-helices, consistent with the flexible architecture required for a multi-epitope construct (**S3 Fig**). The refined tertiary structure of the MEV showed acceptable stereochemical quality and overall structural reliability as shown in **Fig 8**. SWISS-MODEL Ramachandran analysis indicated that 85% of residues were located in favored regions, while PROCHECK showed that 80% of residues fell within the favored regions. Additionally, the ProSA-web Z-score of the construct was found to be –1.47, falling within the acceptable range for native proteins of comparable size.

**Fig 8.**
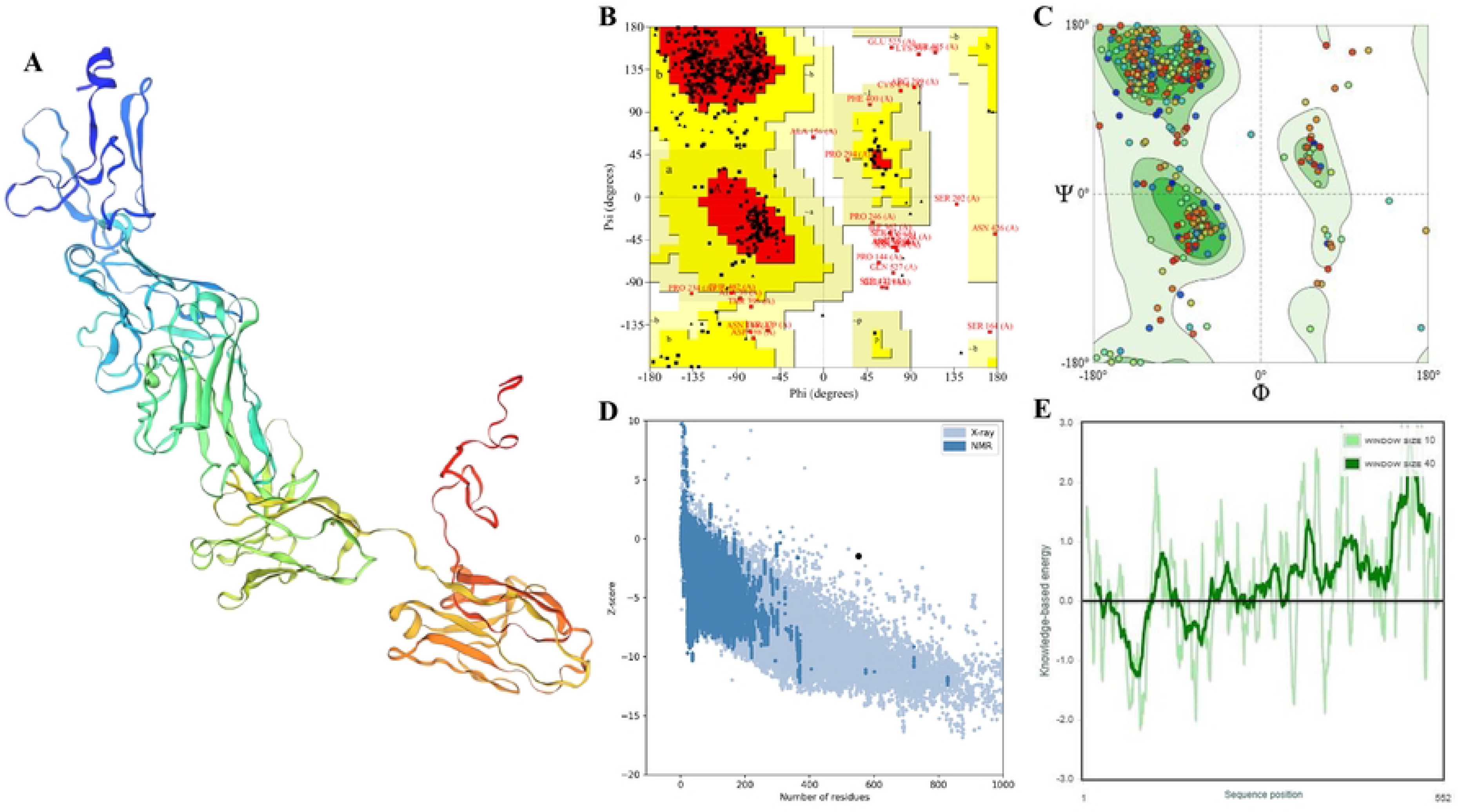
Tertiary structure modelling and assessment of the MEV construct. (**A**) Illustration of the refined 3D structure of the MEV (**B**) PROCHECK Ramachandran plot analysis (**C**) Ramachandran plot analysis by SWISS-MODEL Structure Evaluation (**D**) Z-score of the model relative to experimentally determined structures of similar size through ProSA-web (**E**) ProSA-web residue-wise energy profile of the MEV sequence.

Disulfide engineering recognized 86 intra-chain residue pairs within the MEV model as potential mutation candidates. Among these, 13 residue pairs, namely Ala4-Ala52, Thr110-Gly113, Pro132-Leu148, Ser164-Pro212, Thr198-Pro292, Gly235-Asn250, Pro294-His303, Tyr351-Ala353, Val360-Tyr364, Lys393-Lys401, Gln396-Lys399, Pro429-Thr513 and Phe481-Asp514 satisfied the strict geometric and energetic filters chosen for disulfide bond formation. These residue pairs were therefore retained as the most favorable candidates to enhance the structural stability of the construct (**S4 Fig**). In parallel, six conformational B-cell epitope clusters were mapped onto the refined three-dimensional vaccine model (**S10 Table**), outlining their spatial distribution across the surface of the vaccine (**S5 Fig**).

### MEV-TLR4 interaction analysis, immune simulation, and *in silico* cloning

Molecular docking of the MEV with human TLR4 generated multiple structural complexes, and the top-ranked model corresponded to the lowest-energy cluster with a score of –985.7, as shown in **Fig 9A**. Binding affinity analysis at 37 °C predicted a binding free energy (ΔG) of – 16.4 kcal/mol and a dissociation constant (Kd) of 2.7 x 10^-12^ M, reflecting a highly stable, strong protein-protein interaction interface. Interface analysis using PDBsum revealed two major contact regions between the receptor and the MEV construct (**Fig 9B**). In the primary interface (**Fig 9C**), 32 vaccine residues and 43 receptor residues contributed interface areas of 1,917 Å² and 1,805 Å², respectively, stabilized by several hydrogen bonds. In the second interface (**Fig 9D**), two vaccine residues and six receptor residues contributed interface areas of 222 Å² and 202 Å², respectively, held together by multiple hydrogen bonds. No substantial interactions were detected between the MEV and the remaining TLR4 chains.

**Fig 9.**
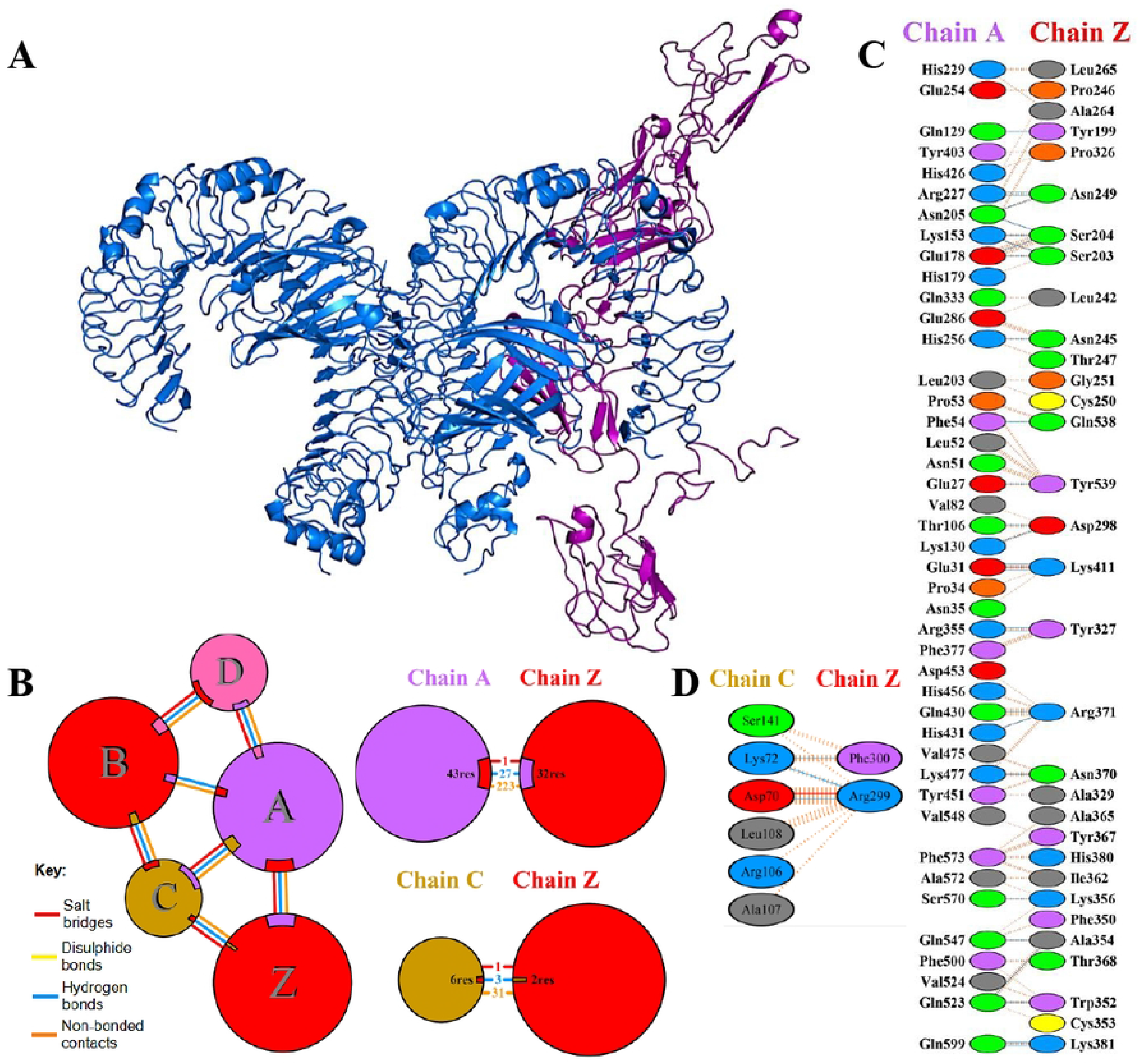
Interface analysis of the MEV-TLR4 complex. (**A**) Docked complex of the MEV (purple) and human TLR4 (blue). (**B**) PDBsum schematic of chain-level contacts, where circle area is proportional to chain surface area and colored lines denote hydrogen bonds (blue), disulfide bonds (yellow), salt bridges (red), and non-bonded contacts (orange). (**C**) Residue-level interactions across the interface between TLR4 (chain A) and vaccine (chain Z). (**D**) Residue-level interactions across the interface between TLR4 (chain C) and vaccine (chain Z).

The intrinsic motions and structural stability of the docked MEV-TLR4 complex were further supported by normal mode molecular dynamics simulation analysis (**Fig 10**). The deformability profile displayed only a few pronounced peaks, whereas most residues displayed low mobility (**Fig 10A**). In agreement with this, the B-factor plot indicated only minor atomic fluctuations along the complex backbone (**Fig 10B**). The first eigenvalue of the dominant normal mode was determined to be 7.894042 × 10^-6^ (**Fig 10C**), while the variance contributed by each individual mode showed the expected inverse relationship with its corresponding eigenvalue (**Fig 10D**). The covariance (**Fig 10E**) and elastic-network analyses (**Fig 10F**) suggested highly coordinated motions and mechanically rigid domains within the core complex. These findings confirm that the MEV-TLR4 complex is structurally stable, mechanically robust, and dynamically coherent.

**Fig 10.**
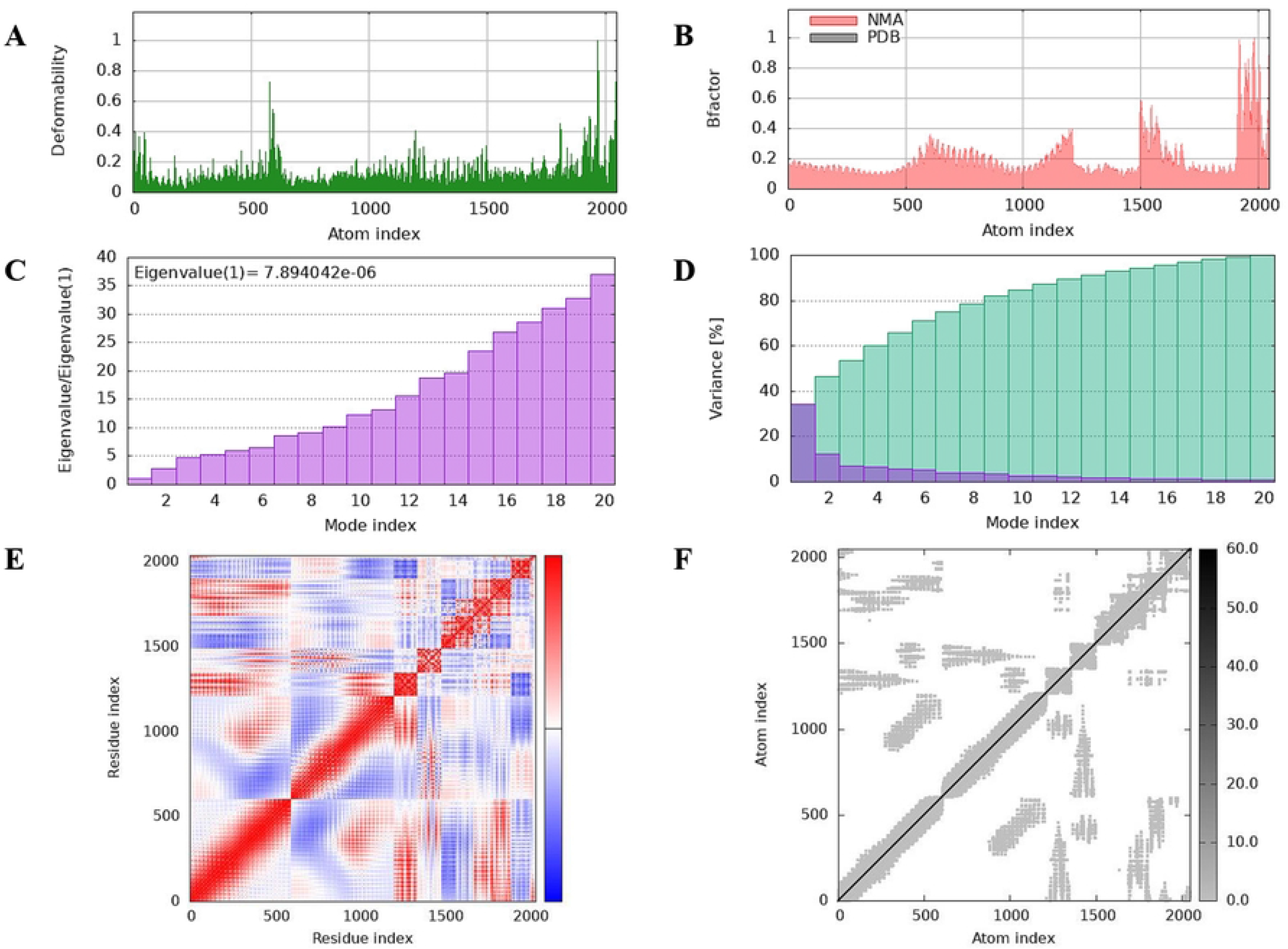
Molecular dynamics simulation analysis of the MEV-TLR4 complex. (**A**) Deformability plot demonstrating flexibility residue-wise. (**B**) B-factor plot comparing predicted B-factors with experimental PDB values. (**C**) Eigenvalue distribution, reflecting the stiffness of the complex, (**D**) Variance associated with each mode (purple) and cumulative variance (green). (**E**) Covariance map of atomic displacements, where correlated, anti-correlated, and uncorrelated motions are indicated by red, blue, and white, respectively. (**F**) Elastic network representation in which darker grey dots indicate stiffer springs.

Immune simulation using C-ImmSim predicted a strong, recallable protective response following three sequential doses of the MEV. Repeated antigen exposure was associated with rapid antigen clearance, expansion of helper and cytotoxic T-cell populations, accumulation of memory cells, and increased B-cell and plasma-cell responses (**Fig 11**). The helper T-cell (T_H_) populations, including memory subsets, expanded sharply after each dose and contracted thereafter, while retaining sizeable long-term memory pools (**Fig 11A**). The T_H_ cell distribution across functional states like resting, active, and duplicating states confirmed cycles of activation and proliferation (**Fig 11B**). The cytotoxic T-cells (T_C_) maintained relatively stable total counts with more modest but sustained responses compared to T_H_ cells, with evidence of repeated activation and persistence of memory cells across the dosing schedule (**Fig 11G**, **Fig 11H**). Regulatory T-cells increased transiently and gradually returned toward baseline (**Fig 11C**). B-cell responses mirrored this pattern, with total and memory B-cell populations increasing over the course of immunization (**Fig 11D**). State-resolved B-cell kinetics showed that B-cells periodically entered active, internalizing and antigen-presenting states, together with transient bursts of plasma B-cells after each dose, indicating efficient differentiation into antibody-secreting cells (**Fig 11E**, **Fig 11F**).

**Fig 11.**
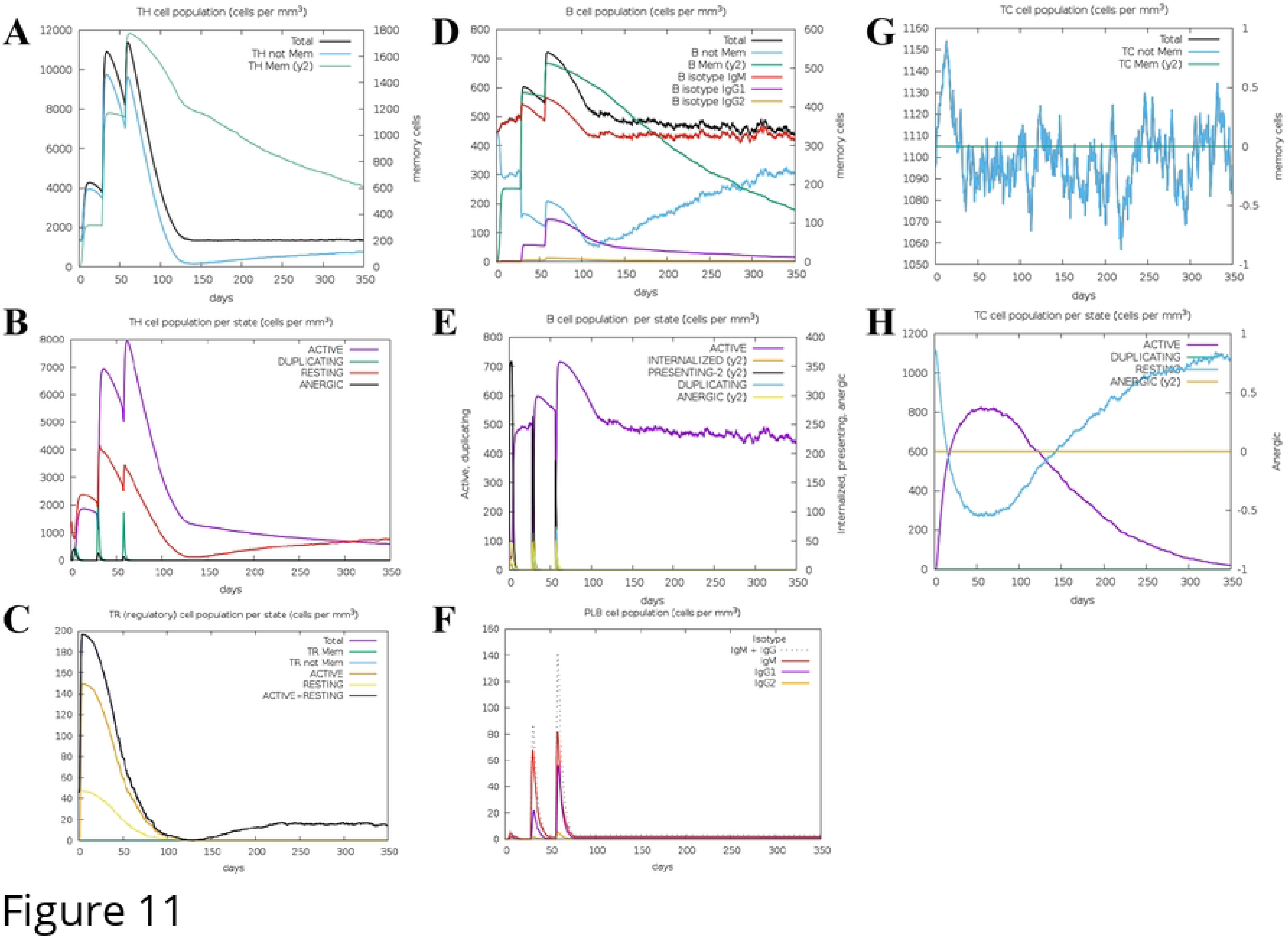
*In silico* immune response profiles elicited by the MEV. (**A**) CD4⁺ T-helper (TH) cell population over time showing total, non-memory and memory subsets. (**B**) TH cells subdivided by state (active, duplicating, resting and anergic). (**C**) CD4⁺ regulatory T (TR) cells, including total, memory and state-specific populations. (**D**) B-cell population showing total and memory cells and class-switched isotypes (IgM, IgG1 and IgG2). (**E**) B cells per functional state (active, internalized antigen, presenting on MHC II, duplicating and anergic). (**F**) Plasma B (PLB) cell population with secreted antibody isotypes IgG1, IgG2, and IgM. (**G**) CD8⁺ cytotoxic T (TC) cell population showing total and memory subsets. (**H**) TC cells subdivided by state (active, duplicating, resting and anergic).

The humoral profile was characterized by an initial IgM-dominated response followed by marked class switching during booster doses, with combined IgG1 and IgG2 levels exceeding IgM as antigen levels declined (**Fig 12A**). Cytokine kinetics revealed sharp, vaccine-driven peaks of IL-2, IFN-γ, and other interleukins after each dose, consistent with repeated activation of T– and B-cell responses and the establishment of long-term immunological memory (**Fig 12B**). In parallel, macrophages (MA) and dendritic cells (DC) maintained antigen-processing and antigen-presenting states while their total populations remained relatively stable (**Fig 12C**, **Fig 12E**). Epithelial cells remained comparatively stable, with only low levels of infected or presenting cells (**Fig 12D**), and NK cells fluctuated moderately around a stable baseline without evidence of excessive activation (**Fig 12F**).

**Fig 12.**
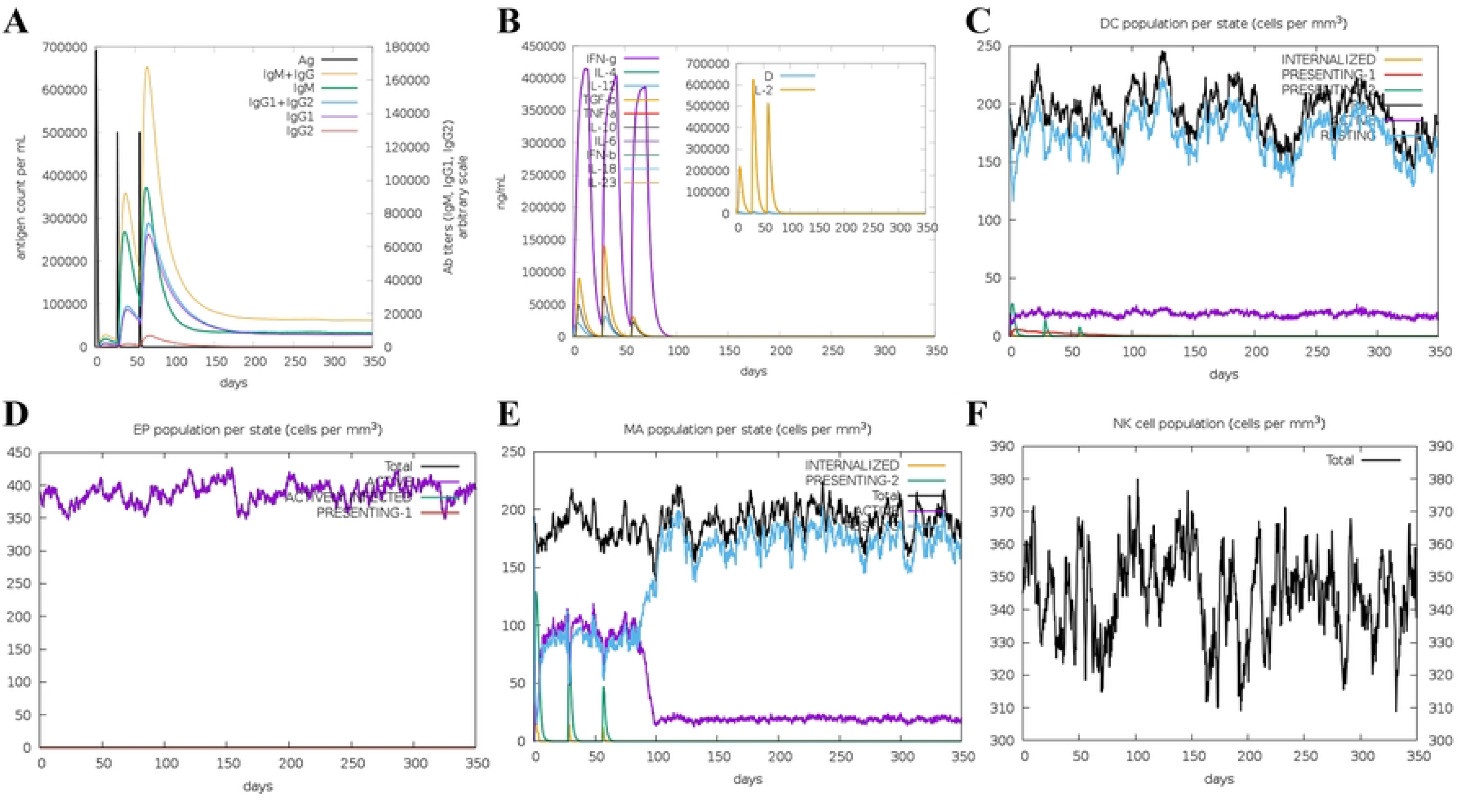
*In silico* humoral and innate immune profiles elicited by the MEV. (**A**) Antigen clearance and antibody titers over time, with antibodies resolved by isotype (IgG1, IgG2, and IgM and their combinations). (**B**) Kinetics of key interleukins and cytokines, and the inset shows the associated danger signal D. (**C**) Dendritic cell (DC) population per state, including total, resting, antigen-internalized and antigen-presenting (MHC class I and II). (**D**) Epithelial cell (EP) population per state, showing total, active, virus-infected and class I–presenting cells. (**E**) Macrophage (MA) population per state, including total, resting, active, internalized and MHC class II–presenting cells. (**F**) Natural killer (NK) cell population (total).

The feasibility of heterologous expression of the MEV in *E. coli* K-12 was further supported by codon optimization and *in silico* cloning. The optimized coding sequence was 1,656 bp in length, with a GC content of 53.7% and a codon adaptation index of 1.00, both consistent with efficient bacterial expression (**S6 Fig**). After incorporation of Eco53kI and BstZ17I restriction sites at the 5′ and 3′ ends, respectively, the optimized sequence was cloned *in silico* into the expression vector to yield a final recombinant plasmid construct, as illustrated by the circular map in **Fig 13**.

**Fig. 13.**
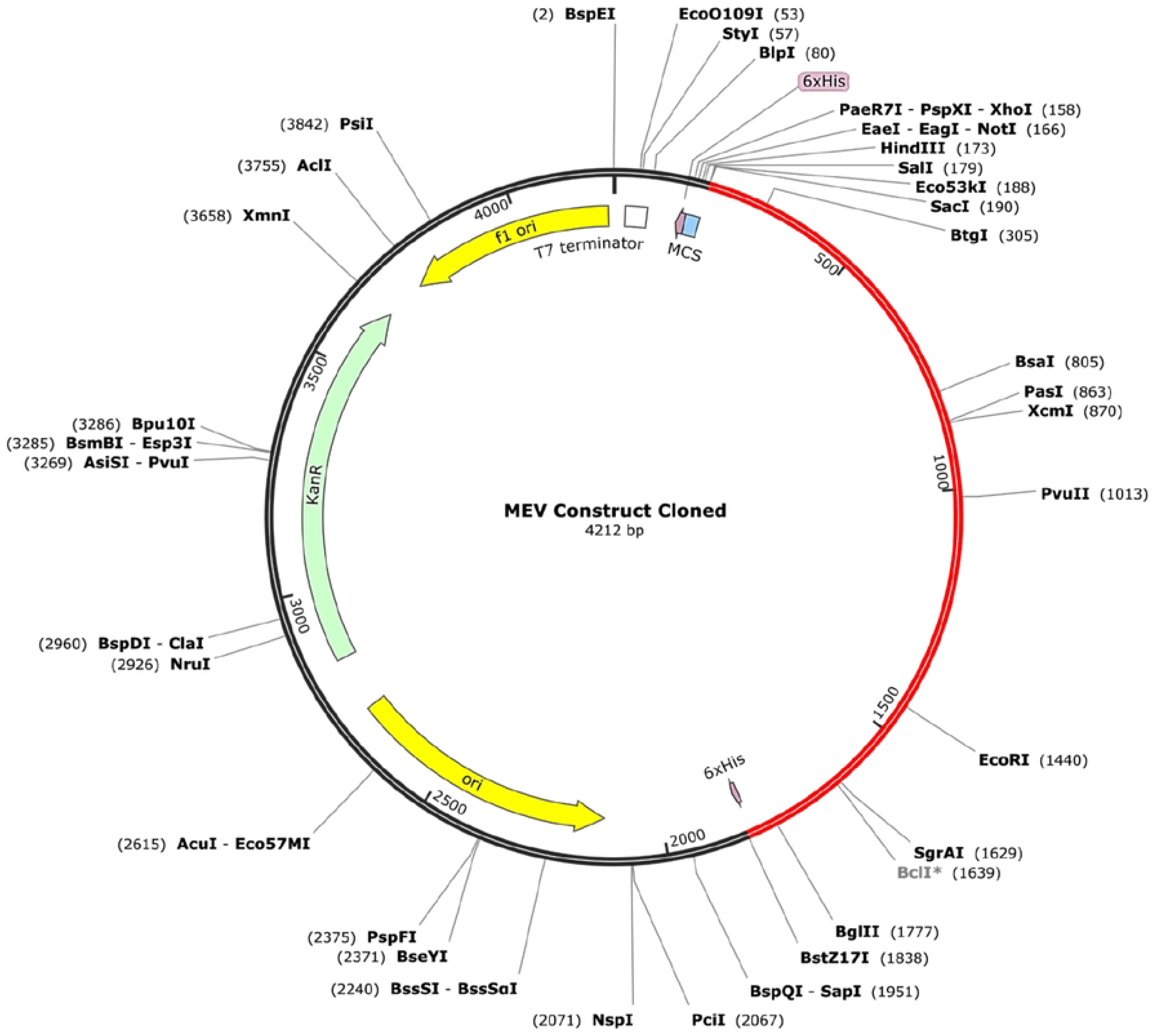
*In silico* cloning of the MEV construct into pET-28a (+). Circular map of the recombinant pET-28a (+) plasmid showing the codon-optimized MEV insert (red) within the vector backbone (black), including the relevant restriction enzyme recognition sites for expression in *E. coli*.

## Discussion

*P. multocida* Pm70 is a non-motile zoonotic coccobacillus that causes severe, life-threatening infections across animals, poultry, and humans [3,7,20,140–142]. Over the last decade, this pathogen has developed resistance to conventional antimicrobial agents, and the absence of an approved human vaccine highlights the urgent need for new therapeutic and preventive interventions [14,15,20,143,144]. In this study, we screened the entire proteome of *P. multocida* Pm70 and identified target proteins that have the potential to serve as both drug and vaccine targets using an integrated pipeline of computational subtractive genomics, reverse vaccinology, and immunoinformatics. Among the prioritized candidates, two outer membrane proteins, OMP47 and LptD, emerged as compelling targets based on stringent filter criteria and comprehensive structural analyses. This was then accompanied by the strategic construction and *in silico* validation of a chimeric MEV construct. Compared to earlier subtractive-genomics and vaccine-design workflows, the present study successfully links therapeutic-target prioritization with vaccine-antigen selection for OMP47 and LptD, providing a promising combination-based strategy for future therapeutic investigations [48–51,145,146].

The prioritized outer membrane proteins, OMP47 and LptD, are compelling dual-purpose therapeutic targets not only because they are surface-exposed proteins but also due to their central, indispensable biological roles in *P. multocida*. The 47 kDa outer membrane protein, or OMP47, is a surface-exposed β-barrel outer-membrane protein that functions in adhesion and transport across the outer membrane region of the pathogen [60–64]. Conversely, the LPS-assembly protein LptD serves as the terminal outer membrane β-barrel component of the LptDE translocon complex and, together with LptE, facilitates the insertion of lipopolysaccharide (LPS) molecules into the outer leaflet of the outer membrane, a process which is central to final LPS and outer membrane assembly [52–59].

In recent years, addressing the increasing antimicrobial resistance among *P. multocida* strains has become a serious concern, as numerous isolates have developed resistance to multiple drug classes including β-lactams, tetracyclines, erythromycin, macrolides, and fluoroquinolones [7,14–17]. Targeting LptD could critically compromise the LPS-assembly of the pathogen and destabilize its outer membrane, while OMP47 inhibition could directly interfere with adhesion– and transport-associated functions. Moreover, our study also indicates that both OMP47 and LptD serve as central hubs within crucial protein-protein networks inside *P. multocida*, interacting with three or more neighboring proteins. Therefore, inhibition of these proteins may disrupt the stability and proper functioning of other associated proteins. Furthermore, the identification of specific binding pockets and representative ligands establishes a structural starting point for the development of novel small-molecule inhibitors that can be designed to disable these target proteins.

As these two proteins are integral to pathogen survival and pathogenesis, we evaluated their potential as viable vaccine targets by examining their antigenicity. Both OMP47 and LptD were predicted to be highly antigenic. Candidate CTL, HTL, and LBC epitopes were prioritized based on stringent selection filters. High-confidence epitopes were assembled along with adjuvants and class-specific linkers in the construction of the chimeric MEV to ensure proper tertiary folding, enhance immunogenicity, and improve structural stability. The designed MEV construct was predicted to be antigenic, non-toxic, non-allergenic, and possessed favorable physicochemical attributes. Structural modelling, refinement and validation further confirmed the acceptable quality of the vaccine construct. Moreover, the strong and stable interaction profiles observed during protein-protein docking, binding affinity calculations, and normal mode analysis simulations indicated that the MEV can effectively engage human TLR4, thereby initiating the innate immune signalling cascades required for a strong vaccine response. Immune simulations further demonstrated its capacity to trigger coordinated innate and adaptive responses, characterized by the increased secretion of cytokines and immunoglobulins, the rapid expansion of B– and T-cell populations, and the formation of immunological memory after each consecutive dose. Furthermore, successful *in silico* cloning into the pET-28a (+) vector optimized for *E. coli* systems indicates that that this vaccine candidate could be efficiently mass-produced using standard bacterial fermentation processes.

Furthermore, a BLASTp analysis using an E-value cutoff of 10^-4^ revealed that both target proteins are conserved across several high-priority pathogens, including *Neisseria gonorrhoeae* and *Pseudomonas aeruginosa*, among others, while demonstrating minimal homology to commensal microbes, indicating their potential to serve as cross-species targets with reduced risks of dysbiosis. LptD, in particular, demonstrated a wider distribution and stronger sequence alignment across species compared to OMP47, while both target proteins revealed little to no similarities with human gut microbiome (**S14 Table**). These results align with the established biological relevance of LptD as an essential protein in outer membrane integrity and virulence of most Gram-negative species such as *E. coli* [147–149]. The translational importance of this target is further highlighted by the antipseudomonal agent murepavadin, which specifically targets LptD in *Pseudomonas aeruginosa* [150]. Furthermore, numerous studies have identified and characterized LptD as a promising vaccine antigen against *Vibrio species* and *Neisseria gonorrhoeae* [151–153]. Similarly, OMP47 homologs observed in species like *Haemophilus influenzae* are known to play critical roles in cellular adhesion and immune recognition [154–156]. While the additional essential proteins identified in this study provide a wider reservoir of targets for future combination-based drug and vaccine design, it requires further in-depth structural and functional investigations, which lies beyond the scope of the current article. Although further experimental validation is recommended to fully assess the *in vitro* and *in vivo* efficacy of these prioritized proteins, the present study establishes a solid foundation for the development of novel therapeutics and vaccine candidates against *P. multocida*, ultimately contributing to the reduction of its rising infections.

## Conclusion

In this study, using an advanced computational pipeline integrating subtractive genomics, reverse vaccinology, and immunoinformatics, we successfully identified two outer membrane proteins, OMP47 and LptD, with integral biological roles in the viability and pathogenesis of *P. multocida*, as promising dual-purpose drug and vaccine targets. Epitope mapping from these two targets further enabled the construction of a viable MEV candidate with favorable structural and physicochemical characteristics. Further investigations revealed that these two targets are well-conserved across several high-priority Gram-negative species beyond *P. multocida*, while displaying minimal similarity to beneficial human gut microbiota. While these findings provide a strong foundation for future antimicrobial and vaccine development against *P. multocida*, further validation through *in vitro* and *in vivo* experiments is recommended to fully assess their efficacy, which remains an essential next step to translate these computational insights into tangible therapeutic solutions.

## Supporting Information

**S1 Fig. ProSA-web assessment of the three-dimensional protein structures of OMP47 and LptD.** (**A**) Overall quality assessment, (**B**) Local quality assessment, (**C**) Lower and higher energy regions of the three-dimensional OMP47 structure. (**D**) Overall quality assessment, (**E**) Local quality assessment, (**F**) Lower and higher energy regions of the three-dimensional LptD structure.

**S2 Fig. Three-dimensional representations of the ligands used for binding-site analysis and docking.** (**A**) ADA or Alpha-D-galactopyranuronic acid, with the IUPAC name (2S,3R,4S,5R,6S)-3,4,5,6-tetrahydroxyoxane-2-carboxylic acid, originated from a *Dickeya dadantii* protein template (PDB ID: 4PR7) and (**B**) C8E or (hydroxyethyloxy)tri(ethyloxy)octane, with the IUPAC name 2-[2-[2-(2-octoxyethoxy) ethoxy]ethoxy]ethanol was derived from a *Shigella flexneri* protein template (PDB ID: 4Q35).

**S3 Fig. Design of the MEV construct.** (**A**) Linear schematic of the MEV showing the N-terminal human β-defensin-3 adjuvant linked by an EAAAK linker, followed by HTL, CTL and LBC epitopes joined by GPGPG, AAY and KK linkers, respectively, and a C-terminal 6xHis-tag (**B**) Full amino acid sequence of the MEV, with the adjuvant in purple, HTL, CTL and LBC epitopes in blue, red and green, respectively, and linkers and His-tag residues in black. (**C**) Predicted secondary structure of the MEV, indicating a prevalence of random coils (grey), with β-strands (yellow) and α-helices (pink).

**S4 Fig. Disulfide engineering of the MEV.** (**A**) Backbone representation of the original model without engineered disulfide bonds (**B**) Mutant model after disulfide engineering, showing 13 introduced disulfide bridges (yellow) between selected residue pairs that satisfy standard disulfide geometry and energy criteria.

**S5 Fig. Conformational B-cell epitope clusters on the three-dimensional MEV structure.** Panels (**A-F**) show individual conformational B-cell epitope clusters (yellow surface) mapped onto the vaccine model (grey sticks).

**S6 Fig. *In silico* cloning map and GC content profile of the MEV construct.** (**A**) Linear representation of the codon-optimized MEV sequence showing restriction endonuclease recognition sites. (**B**) GC content distribution across the MEV construct, with color-coded bars indicating local GC percentage along the sequence.

**S1 Table. Proteins essential for the survival of *P. multocida*.** This lists all 129 essential proteins found to be crucial for the survival of *P. multocida*.

**S2 Table. Essential proteins involved in important pathogen-specific metabolic pathways with no homology with known drug targets in the DrugBank database.** This lists the metabolic pathways with their KEGG pathway IDs, the number of proteins per pathway and their corresponding UniProt accession numbers.

**S3 Table. Pathway-independent proteins with no homology with known drug targets in the DrugBank database.** This lists the proteins with their UniProt IDs, gene names, protein names, available drugs, and predicted biological functions.

**S4 Table. Essential proteins associated with virulence.** This lists the virulent proteins identified by VirulentPred2.0 along with their functions, UniProt ID and KO ID.

**S5 Table. The subcellular localization of all shortlisted essential proteins.** This lists the subcellular localization of all shortlisted essential proteins identified by pSORTb v3.03 and CELLO v2.5.

**S6 Table. The list of proteins interacting with the two target proteins, OMP47 and LptD.** This lists the details of interacting proteins along with their STRING scores.

**S7 Table. Detailed analyses on the binding sites of both proteins, OMP47 and LptD**. This lists the characteristics, size, shape, functional group, amino acid descriptors and compositions of the top-ranked binding sites of both target proteins.

**S8 Table. Amino acid residues forming the top-ranked binding pockets of OMP47 and LptD**. This lists the corresponding chains and residues that constitute the binding sites and their sequence positions.

**S9 Table. Physicochemical and immunological properties of the MEV construct.** The physicochemical and immunological features of the MEV are listed.

**S10 Table. Predicted conformational B-cell epitope clusters within the MEV construct**. For each cluster, the constituent residues, number of residues, and corresponding ElliPro score are listed.

**S11 Table. All CTL epitopes identified from both target proteins, OMP47 and LptD.** This lists all CTL epitopes with immunogenicity scores, antigenicity values, predicted toxicity and allergenicity profiles, percentile ranks, prediction scores, and peptide lengths.

**S12 Table. All HTL epitopes identified from both target proteins, OMP47 and LptD.** This lists all HTL epitopes with antigenicity scores, predicted toxicity and allergenicity profiles, percentile ranks, prediction scores, peptide lengths, and cytokine inducing potentials.

**S13 Table. All LBC epitopes identified from both target proteins, OMP47 and LptD.** This lists all LBC epitopes with peptide lengths, antigenicity scores, predicted toxicity and allergenicity profiles, transmembrane topology, and their capacity to induce IgG, IgM, and IgA responses.

**S14 Table. Cross-species BLASTp.** This lists the cross-species BLASTp analysis of the two proteins, OMP47 and LptD, against high-priority pathogens and commensal microbes indicating their potential as cross-species targets.

